# Post-transcriptional regulation by TIA1 and TIAL1 controls the transcriptional program enforcing T cell quiescence

**DOI:** 10.1101/2024.09.03.608755

**Authors:** Ines C. Osma-Garcia, Orlane Maloudi, Mailys Mouysset, Dunja Capitan-Sobrino, Trang-My M. Nguyen, Yann Aubert, Manuel D. Diaz-Muñoz

**Affiliations:** Toulouse Institute for Infectious and Inflammatory Diseases (INFINITy), Inserm UMR1291, CNRS UMR5051, University Paul Sabatier, CHU Purpan, Toulouse, 31024, France

**Keywords:** T cells, homeostasis, quiescence, post-transcriptional gene regulation, RNA binding proteins TIA1, TIAL1, FOXP1, transcriptional networks

## Abstract

Immune protection against new and recurrent infections relies on long-term maintenance of a highly diversified T-cell repertoire. Transcription factors cooperate to enforce T-cell metabolic quiescence and maintenance. However, less is known about the post-transcriptional networks that preserve peripheral naïve T cells. Here we describe the RNA binding proteins TIA1 and TIAL1 as key promoters of CD4 and CD8 T cell quiescence. T cells deficient in TIA1 and TIAL1 undergo uncontrolled cell proliferation in the absence of cognate antigens, leading this to a premature T-cell activation, exhaustion and death. Mechanistically, TIA1 and TIAL1 control the expression of master regulatory transcription factors, FOXP1, LEF1 and TCF1, that restrain homeostatic T-cell proliferation. In summary, our study highlights a previously unrecognised dependency on post-transcriptional gene regulation by TIA1 and TIAL1 for implementing the quiescent transcriptional programs for long survival of T cells.

## INTRODUCTION

Immune system evolution primes development of T cells with an ample repertoire of T-cell receptors (TCRs) to fight pathogens and foreign antigens. The number of naïve T cell clones kept in peripheral organs is over 10^9^ ^1^, with only a handful of clonotypes detecting a given foreign antigen to rapidly mount an immune response. In the thymus, rearrangement of the TCR locus enables generation of this highly diversified T cell pool for which mild recognition of MHC-restricted antigen receptors loaded with self-peptides (self-pMHC) drives T cell selection and maturation ^2, 3^. In the periphery, TCR:self-pMHC interaction provides a tonic signal for long-term maintenance of naïve T cells in quiescence until a cognate antigen is detected.

Quiescence is the cellular state of low metabolic activity and cell cycle arrest. Signalling from TCR:self-pMHC and checkpoint receptors (e.g. VISTA, CTLA-4) enforce specific transcriptional programs for the entry and maintenance of T cells in quiescence ^4^. Several transcription factors (TFs) cooperate on the establishment of this quiescent program, including forkhead box (FOX) proteins, basic region-leucine zipper (bZip), Kruppel like factors (KLFs), and high mobility group (HMG) family members ^5, 6, 7, 8^. They act as transcriptional gene repressors of the energy metabolism and cell cycle, and sustain the expression of the pro-survival factor BCL2 ^9, 10^. Quiescence-related TFs are characterised by having a relative short protein half-life ^11^ that, along with active post-transcriptional regulation of their messenger (m)RNA stability ^12, 13, 14^, enables maintenance of T cells in a state of preparedness for rapidly turning on the proliferative and terminal differentiation programs that produce abundant T-helper cells. Despite this knowledge, the interconnection between transcriptional and post-transcriptional programs in quiescent T cells remains largely unexplored.

RNA and protein abundance poorly correlate in T cells (r^2^=0.4) ^11, 15, 16^. This highlights the existence of multiple layers of post-transcriptional gene regulation that control T cell quiescence and activation. Indeed, foundational studies on the modulation of cytokine production (e.g. TNF, IFNγ,…) revealed evolutionary conserved RNA regulatory programs responsible for the discrepancies between mRNA and protein expression ^17, 18, 19^. RNA splicing, editing, transport, location, translation and decay shape the production of proteins in cells, both quantitatively and qualitatively ^20^. The number of RNA transcripts annotated in lymphocytes is more than three times higher the number of genes due to alternative splicing (AS) (67,453 RNA transcripts versus 20,766 genes) ^21^. Genetic information is further amplified by alternative translation of mRNAs due to differential usage of start codons, reading frame switch and the presence of internal ribosome entry sites (IRES) ^22, 23^. In addition, recent approaches combining transcriptomics, ribosome profiling, mass spectrometry and computational analysis have enabled the identification of thousands of new open reading frames (ORFs) in lymphocytes with yet unknown functions ^24, 25^.

RNA binding proteins (RBPs) constitute highly dynamic post-transcriptional hubs during T-cell development and function ^20, 26^. They control, in time and magnitude, the expression of key modulators of lymphocyte quiescence such as FOXP1 and BCL2 ^27, 28^. In addition, RBPs are instrumental for mRNA poising in resting T cells and timely cytokine production upon immune-memory reactivation ^29, 30^. In this study, we describe the T-cell antigen 1 (TIA1) and its paralog TIA like(L)-1 as essential RBPs for the expression of quiescence-enforcing TFs and the long-term maintenance of naïve T cells.

TIA1 was first discovered in cytotoxic CD8 T cells ^31^ and, since then, it has been adopted as a good prognosis marker of anti-tumoral CD8 T cell infiltration in cancer ^32^. Despite this observation, the function of TIA1 and TIAL1 has never being investigated in T cells. TIA1 and TIAL1 are evolutionary conserved proteins, with 77% amino acid sequence homology between paralogs and over 90% conservation between human and mouse. TIA1 and TIAL1 have been extensively studied in cancer cell lines in which, depending on their level of expression and type of cancer, they exert a dual role as oncogenes or tumour suppressor genes ^33, 34, 35^. Changes in the expression of TIA1 are also linked with the development of neuropathies, including Alzheimer’s disease (AD), amyotrophic lateral sclerosis (ALS) and frontotemporal dementia (FTD) ^36, 37, 38^. TIA1 and TIAL1 are essential for embryonic cell development and differentiation. TIAL1 knockout (KO) mice are embryonic lethal ^39^, whereas 50% of TIA1-KO mice die between E16.5 and post-natal week 3. The survivors have profound immunological defects associated to exacerbated production of cytokines (IL6, TNF) in macrophages ^40^. Binding of TIA1 and TIAL1 modulates the mRNA splicing and translation of multiple immune modulators such as p53 ^41^, Fas ^42^, and c-Myc ^43^. Several studies highlight the functional redundancy of TIA1 and TIAL1 in the control of stress responses and the cell cycle ^44^. Recently, we showed that TIA1 and TIAL1 are both required for timely activation of the DNA damage repair machinery during B cell development and selection of antigen-specific B cell clones during vaccination ^45, 46^.

Here we demonstrate that TIA1 and TIAL1 are dispensable during T-cell development, but essential for long-term maintenance of peripheral T cells. Mechanistically, they bind to U-rich elements present in the 3’UTRs of the quiescence-associated TFs FOXP1, LEF1 and TCF1, securing their expression and contributing to the establishment of the transcriptional programs that restrain T-cell activation and differentiation.

## RESULTS

### TIA1 and TIAL1 are required for maintenance of peripheral T cells

Recent proteomics studies have revealed that the expression of TIA1 and TIAL1 is actively modulated in T cells upon TCR engagement ^47, 48^. A closer look to protein copy numbers in both naïve CD4 and CD8 T cells showed that TIAL1 is 4-fold more abundant than TIA1 (**Fig. 1a**). Stimulation of CD4 and CD8 T cells with αCD3 and αCD28 increased the expression of both TIA1 and TIAL1 (**Fig. 1a and Suppl. Fig. 1a**). While the expression of TIA1 augmented by 2- to 4-fold, depending on the cell type and the methodology used for quantitation, the protein levels of TIAL1 increased between 5- to 8-fold after T-cell activation. Further analyses on the expression of TIA1 and TIAL1 in memory-phenotype (MP) and effector-phenotype (EP) T cells revealed that these antigen-experienced cells expressed significantly higher levels of TIA1 and TIAL1 than naïve T cells (**Fig. 1b**), suggesting that these proteins could be important for the establishment of the T cell compartment and/or after activation.

**Figure 1.**
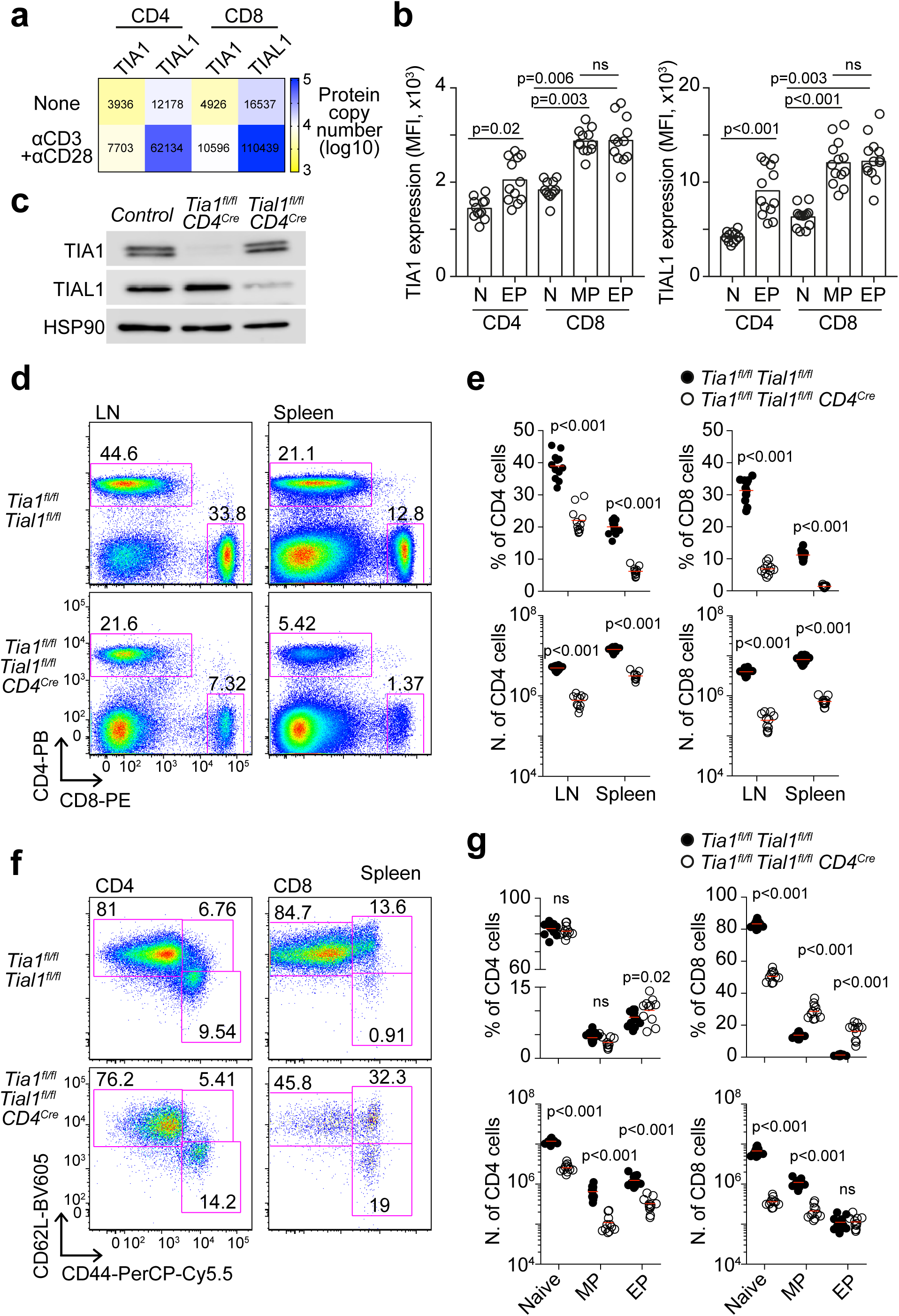
TIA1 and TIAL1 are required for peripheral T cell maintenance. **a,** Heatmap showing TIA1 and TIAL1 protein copy numbers in naive CD4 and CD8 T cells stimulated or not with αCD3 and αCD28 (The Immunological Proteome Resource, ImmPRes ^47, 48^, numbers refer to the mean protein copy number annotated). **b,** Quantitation by flow cytometry of TIA1 and TIAL1 expression in splenic naïve (CD62L^+^ CD44^-^), memory-phenotype (MP, CD62L^+^ CD44^+^) and effector-phenotype (EP, CD62L^-^ CD44^+^) CD4 and CD8 T cells. **c,** Immunoblotting detection of TIA1 and TIAL1 in CD4^+^ thymocytes from control, *Tia1^fl/fl^ CD4^Cre^* and *Tial1^fl/fl^ CD4^Cre^* mice. **d,** Pseudo colour dot plots showing the percentage of CD4 and CD8 T cells in the lymph nodes (LNs) and spleen of control and *Tia1^fl/fl^ Tial1^fl/fl^ CD4^Cre^* mice. **e,** Quantitation of the percentage and number of CD4 and CD8 T cells shown in d. **f,** Pseudo colour dot plots showing naïve, MP and EP T cells in the spleen of control and *Tia1^fl/fl^ Tial1^fl/fl^ CD4^Cre^* mice. **g,** Quantitation of the percentage and the number of naïve, MP and EP T cells. Data in a to g are from at least of two independent experiments. A minimum of 6 mice were used per experiment and genotype in a, e and g. Mann-Whitney tests were performed for statistical analysis.

To uncover the relevance of these proteins in T cells, we generated T-cell specific conditional KO mice by crossing *Tia1^fl/fl^*and *Tial1^fl/fl^* mice with *CD4^Cre^* mice [hereafter called as *Tia1^fl/fl^ Tial1^fl/fl^ CD4^Cre^*mice]. Analysis of Cre-mediated activity in thymic T cell progenitors revealed the efficient recombination of *Tia1* and *Tial1* alleles, with protein depletion achieved from the double positive (DP) stage of thymic T-cell development (**Fig. 1c and Suppl. Fig. 1b-c**), making our model suitable for studying the intrinsic roles of TIA1 and TIAL1 in T cells.

First analyses revealed a significant reduction in the percentage and number of T cells found in the lymphoid organs of *Tia1^fl/fl^ Tial1^fl/fl^ CD4^Cre^* mice compared to control mice (**Fig. 1d and e)**. CD4 T cells were diminished between 2- to 5-fold in the lymph nodes (LNs) and spleen of *Tia1^fl/fl^ Tial1^fl/fl^ CD4^Cre^* mice. Similarly, CD8 T cells were decreased by 10-fold in these mice when compared to control mice. Importantly, expression of an OTII TCR-transgene did not rescue, but rather increased, the loss of CD4 T cells in the absence of TIA1 and TIAL1, suggesting that the loss of peripheral T cells is independent of the specificity of the TCR (**Suppl. Fig. 1d**). Finally, single conditional deletion of *Tia1* or *Tial1* did not alter the percentage or the number of T cells in peripheral lymphoid organs (**Suppl. Fig. 1d, 1e and 1f**) suggesting that, despite the differences in overall protein abundance, TIA1 and TIAL1 act redundantly in the maintenance of T cells.

Phenotypic characterization of T cells in the spleen based on the expression of the homing receptor CD62L and the activation marker CD44 showed a reduction in the proportion of naïve T cells and a concomitant increase in the proportion of MP and EP T cells in *Tia1^fl/fl^ Tial1^fl/fl^ CD4^Cre^* mice (**Fig. 1f and 1g**). This was remarkably significant in CD8 T cells, for which the percentage of EP CD8 T cells increased more than 10-fold in *Tia1^fl/fl^ Tial1^fl/fl^ CD4^Cre^* mice when compared to control mice. Similar data was collected from peripheral LNs (data not shown). Taken together, our data reveal that the expression of TIA1 and TIAL1 changes between T cell subsets and they are required for maintenance of CD4 and CD8 T cells in the periphery.

### TIA1 and TIAL1 are dispensable during thymic development

T-cell lymphopenia in *Tia1^fl/fl^ Tial1^fl/fl^ CD4^Cre^* mice could have its origin in early defects during thymic development. Indeed, TIA1 and TIAL1 are key components of the splicing machinery that controls the synthesis of DNA damage sensors and repair proteins for VDJ gene recombination ^45^. Thus, we tested the hypothesis of whether TIA1 and TIAL1 were needed during early T-cell development in the thymus. First, using flow cytometry, we observed that the expression of TIA1 and TIAL1 was higher in double negative (DN) CD4 and CD8 thymocytes, but drastically diminished upon differentiation into double positive (DP) T cells (**Fig. 2a**). Surface expression of the TCR and activation of DP T cells correlated with a moderate, but sustained, increase in the expression of TIA1 and TIAL1 which remained mostly constant during thymic maturation of single positive CD4 (SP4) and single positive CD8 (SP8) T cells (**Fig. 2a**). Altogether, changes in the expression of TIA1 and TIAL1 at different stages of T-cell thymic development suggested that these RBPs could be involved in the early establishment of the T-cell compartment.

**Figure 2.**
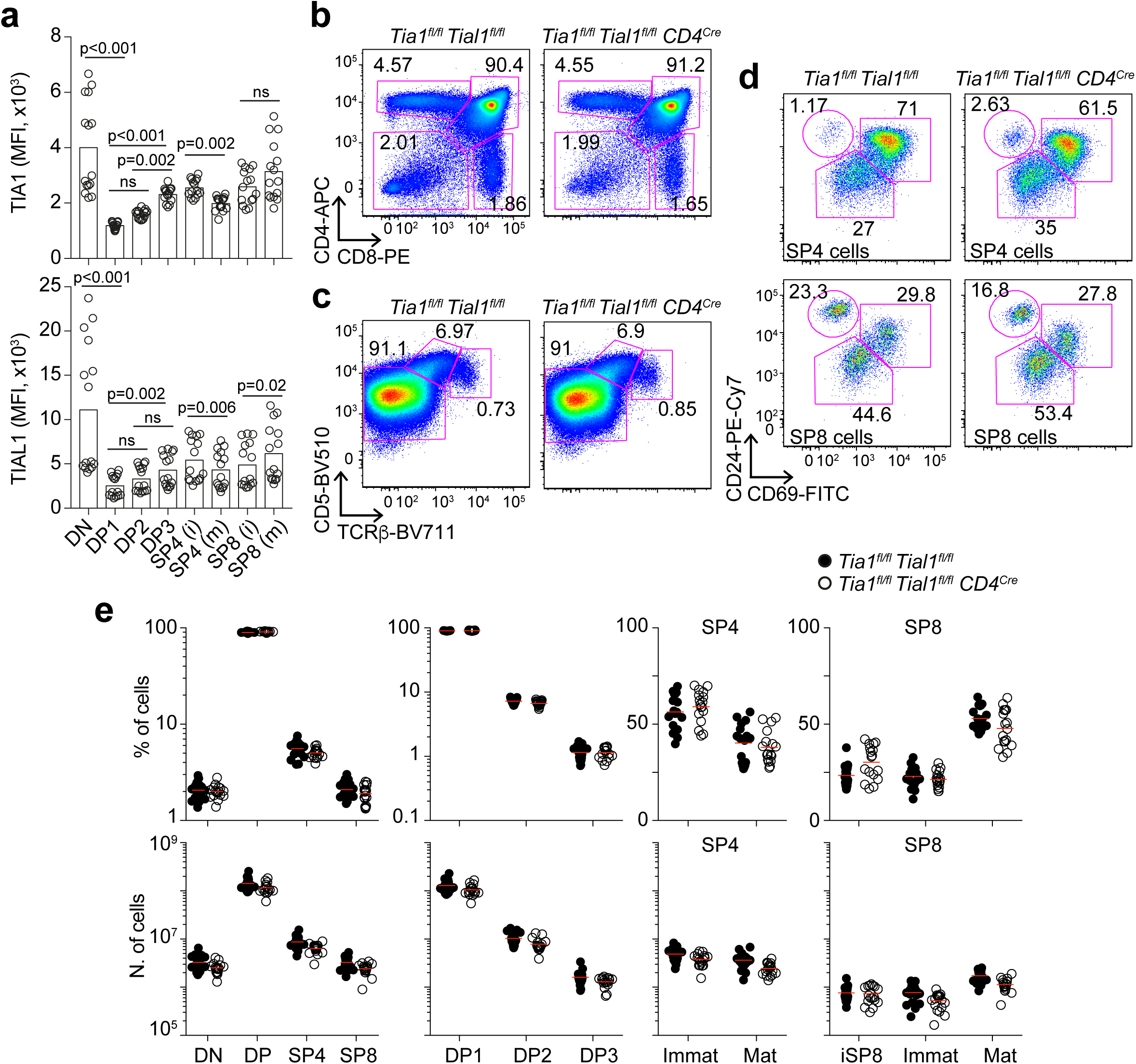
TIA1 and TIAL1 are dispensable during thymic T-cell development. **a,** Quantitation by flow cytometry of TIA1 and TIAL1 expression in thymic T cell subsets. Data pooled from two independent experiments performed each with at least 6 mice. **b,** Representative pseudo-colour dot plots showing the percentage of DN, DP, SP4 and SP8 T cells in control and *Tia1^fl/fl^ Tial1^fl/fl^ CD4^Cre^* mice. **c,** Flow cytometry analysis of DP T cell subsets (DP1, TCRβ^-^ CD5^-^; DP2, TCRβ^int^ CD5^+^; DP3, TCRβ^+^ CD5^+^). **d,** Analysis of thymic SP4 and SP8 cell maturation (immature SP4 and SP8, CD24^hi^ CD69^hi^; mature SP4 and SP8, CD24^lo^ CD69^lo^; iSP8, CD24^hi^ CD69^-^). **e,** Quantitation of T-cell subsets in the thymus of control and *Tia1^fl/fl^ Tial1^fl/fl^ CD4^Cre^* mice. Percentage and number of T cells are from three independent experiments performed each with at least 5 mice per genotype. Mann-Whitney tests were performed in a.

Phenotypic characterization of thymic T-cell subsets in control and *Tia1^fl/fl^ Tial1^fl/fl^ CD4^Cre^* mice revealed no differences in the development of DN to DP T cells (**Fig. 2b and Suppl. Fig. 2**), in the induction of a functional TCR, in positive selection of DP T cells (**Fig. 2c and Suppl. Fig. 2**) or in terminal maturation of SP4 and SP8 T cells (**Fig. 2d and Suppl. Fig. 2**). Therefore, we conclude that, despite the changes in protein expression in thymocytes, TIA1 and TIAL1 are dispensable for thymic development of T cells from the DP stage.

### TIA1 and TIAL1 promote T-cell quiescence

An alternative that could explain the low numbers of peripheral T cells in *Tia1^fl/fl^ Tial1^fl/fl^ CD4^Cre^* mice was that newly produced T cells failed to stop their metabolism to enter into quiescence. Indeed, in the absence of external antigens, most naïve T cells remain at the G0 phase of the cell cycle, with only few cells proliferating in order to maintain the T cell pool. Measurement of homeostatic T cell proliferation in LNs and spleens using the protein Ki67, that labels cell cycle entry into the G1 phase, confirmed the low rate of peripheral T cell self-replenishment in wild-type mice as previously reported ^49^. Interestingly, depending on the T-cell subset, we observed an up to 2-fold increase in the percentage of Ki67^+^ T cells in *Tia1^fl/fl^ Tial1^fl/fl^ CD4^Cre^* mice compared to control mice (**Fig. 3a**). This correlated with a significant increase in the expression of activation and exhaustion markers such as CD25 and CD28, PD1 and CTLA4 in both naïve and antigen-experience T cells (**Fig. 3b**); and with an increment in the percentage of TIA1 and TIAL1 double KO (dKO) EP T cells expressing KLRG1, a marker of exhaustion in highly activated T cells (**Fig. 3c**). Remarkably, naïve CD4 and CD8 T cells were also abnormally activated and exited quiescence in the absence of TIA1 and TIAL1; although, comparatively, to a lower extent than EP T cells (**Fig. 3a and 3b**). Altogether, TIA1 TIAL1 KO T cells show an active and proliferative phenotype which could be related to a loss of quiescence.

**Figure 3.**
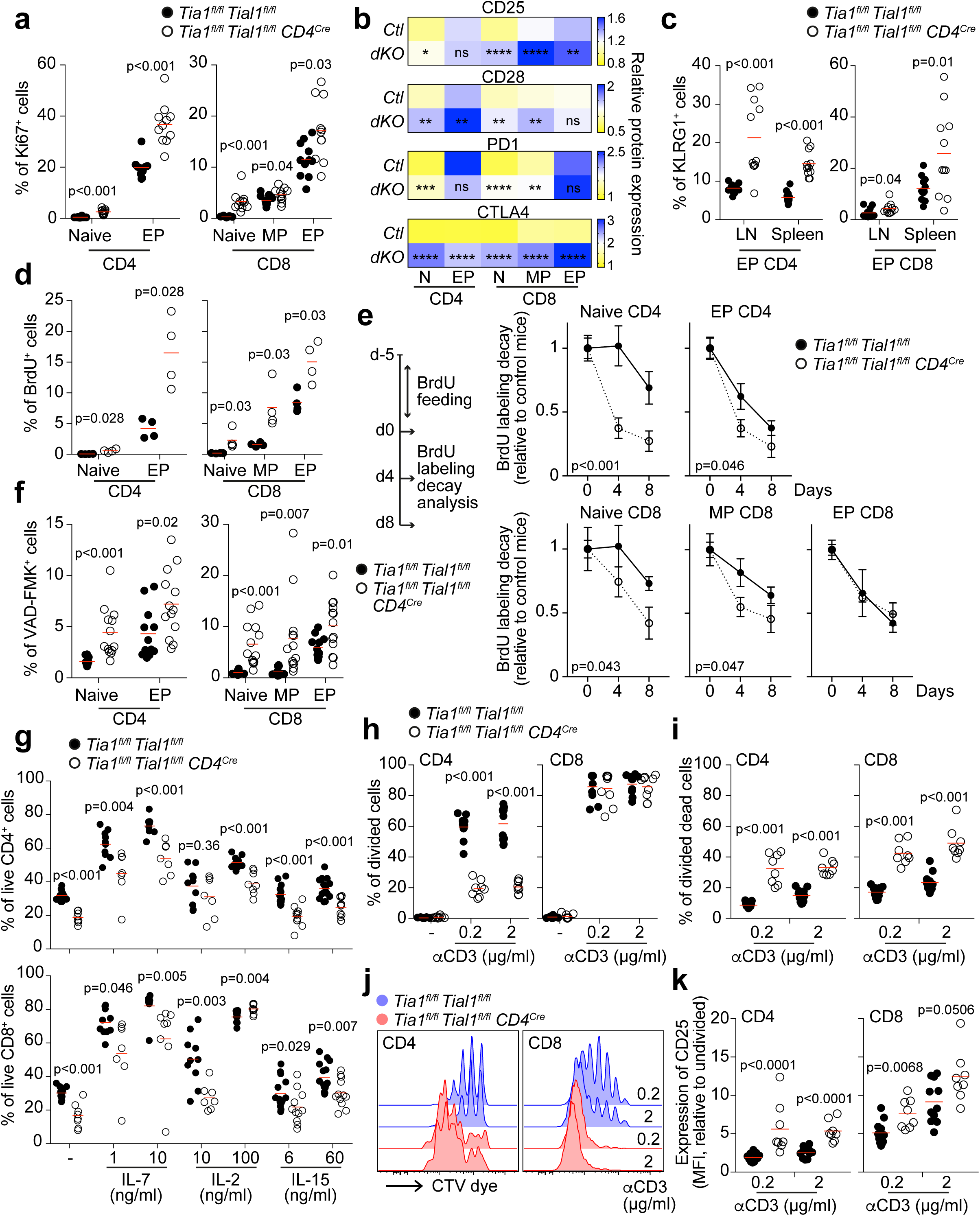
TIA1 and TIAL1 enforce T-cell quiescence. **a,** Analysis of Ki67^+^ LN T cells in control and *Tia1^fl/fl^ Tial1^fl/fl^ CD4^Cre^* mice. **b,** Heatmaps showing the protein surface expression of CD25, CD28, PD-1 and CTLA4 measured in T cells by flow cytometry. Bar scale indicates mean fluorescence intensity (MFI) relative to the expression in naïve CD4 or CD8 T cells. Data is from one or two representative experiment with at least 4 mice per genotype. Mann-Whitney analyses were performed to compare protein expression between similar cell types from from control (Ctl) or *Tia1^fl/fl^ Tial1^fl/fl^ CD4^Cre^* (dKO) mice (* p<0.05, ** p<0.01, ** p<0.001, **** p<0.0001). **c,** Quantitation of the percentage of KLRG1^+^ EP CD4 and CD8 T cells in LNs and spleens of control and *Tia1^fl/fl^ Tial1^fl/fl^ CD4^Cre^*mice. **d,** Measurement of BrdU incorporation into CD4 and CD8 T cells after mouse feeding with BrdU for 5 days. **e,** Analysis of BrdU- labelling decay in CD4 and CD8 T cells from control and *Tia1^fl/fl^ Tial1^fl/fl^ CD4^Cre^* mice at the indicated days after discontinuing BrdU administration. Data shown as relative to the labelling of T cells at day 0 (mean + SEM). **f,** Proportion of CD4 and CD8 T cells with active caspases labelled with FITC-VAD-FMK dye. **g,** Quantitation of the percentage of splenic CD4 and CD8 T cells alive after 3 days *in-vitro* culture in the presence of the indicated concentrations of IL-, IL-7 or IL-15. **h**, Analysis of the total proportion of control and TIA1 TIAL1 dKO CD4 and CD8 T cells that have divided at least once after stimulation with the indicated doses of αCD3. **i,** Measurement of divided dead cells in cell cultures shown in h. **j,** Representative histogram showing cell tracer violet (CTV) dye dilution in proliferating CD4 and CD8 T cells from h. **k,** Expression of the cell surface marker CD25 in CD4 and CD8 T cells proliferating after TCR stimulation. In a and f, data pooled from three independent experiments performed each with at least 4 mice per genotype. In c, e, g, h, I and k, data from two independent experiments performed with at least 4 mice per genotype. Data in h is representative from one of the two independent experiments performed. Two-tailed Mann-Whitney analyses were performed in a, c, d, f, g, h, i and k. Two-way ANOVA was performed in e.

### TIA1 and TIAL1 limit T-cell activation, proliferation and cell death

Next, we combined different *in-vivo* and *in-vitro* approaches to evaluate the activation, proliferation and survival capacity of TIA1 TIAL1 KO T cells. First, we fed control and *Tia1^fl/fl^ Tial1^fl/fl^ CD4^Cre^* mice with BrdU for 5 days and measured BrdU incorporation into the DNA of proliferating CD4 and CD8 T cells. In both spleen and LNs, we detected a higher percentage of BrdU^+^ naïve and EP T cells in *Tia1^fl/fl^ Tial1^fl/fl^ CD4^Cre^* mice (**Fig. 3d**), confirming the increase in homeostatic proliferation in the absence of TIA1 and TIAL1. Further analysis of the decay of BrdU-incorporated DNA after pulse-chase labelling of proliferating T cells revealed a faster reduction in the percentage of BrdU^+^ naïve and EP CD4 T cells, and naïve and MP CD8 T cells from *Tia1^fl/fl^ Tial1^fl/fl^ CD4^Cre^* mice compared to control mice (**Fig. 3e**). Differences in BrdU-labelling decay between control and TIA1 TIAL1 dKO T cells were higher when analysing naïve CD4 and CD8 T cell populations and decreased progressively in MP and EP T cell subset. This was probably a consequence of the higher proliferation capacity of these cells, and remarks further the anomalous entry into cycling of naïve T cells in the absence of TIA1 and TIAL1.

The rapid loss of BrdU-labelled naïve T cells in the absence of TIA1 and TIAL1 could be circumscribed to a rapid turn-over of proliferating cells, to a higher rate of cell death or both. To test these possibilities, we first measured apoptosis by short-term labelling of apoptotic cells with the pan-caspase inhibitor VAD-FMK coupled to FITC and flow cytometry. In line with the overall reduction in CD4 and CD8 T cell populations, we observed an increase in the percentage of apoptotic T cells in *Tia1^fl/fl^ Tial1^fl/fl^ CD4^Cre^* mice compared to control mice. The loss of cell viability ranged from a 1.5-fold to a 4-fold increase depending on the T-cell subset analysed (**Fig. 3f**), and it was independent of changes in cellular energy metabolism (**Suppl. Fig. 2a and 2b**) or protein synthesis (**Suppl. Fig. 2c**).

Decreased survival of TIA1 TIAL1 dKO T cells was further confirmed in *in-vitro* cultures in which CD4 and CD8 T cells were treated with the pro-survival cytokines IL-2, IL-7 or IL-15 (**Fig. 3g and Suppl. Fig. 2d**). Remarkably, this defect was not due alterations in IL-7-mediated signalling and phosphorylation of STAT5 (**Suppl. Fig. 2e**), but most likely associated to the high activation of the TIA1 TIAL1 dKO T cells, which increased the expression of CD44 even in the absence of cytokine (**Suppl. Fig. 2f**). To support this notion, we assessed whether TIA1 TIAL1 dKO T cell survival, activation and proliferation were also altered in response to TCR stimulation. Viability of TIA1 TIAL1 dKO CD4 T cells was highly impaired after 3 days stimulation with anti-CD3 (**Suppl. Fig. 3g**), with only 20% of total (dead or alive) cells undergoing cell division compared to 60% of control cells (**Fig. 3h**). Viability of CD8 T cells was also reduced in the absence of TIA1 and TIAL1, but to a lesser extent than for CD4 T cells (**Suppl. Fig. 2g**). Interestingly, like control CD8 T cells, most of the total TIA1 TIAL1 dKO CD8 T cells underwent proliferation after TCR stimulation. However, up to 50% of these CD8 cells died after dividing at least one time (**Fig. 3i**). This was a 2-fold and a 1.5-fold higher proportion than the percentage of control CD8 T cells and TIA1 TIAL1 dKO CD4 T cells dying after dividing, respectively. Further analysis of the dilution of the Cell Tracer Violet (CTV) dye upon cell division revealed a faster proliferation of CD8 T cells compared to CD4 T cells (**Fig. 3j**). Importantly, both TIA1 TIAL1 CD4 and CD8 T cells were hyperactivated, as reflected by the exacerbated expression of the activation marker CD25 (**Fig. 3k**), and they proliferated more extensively than control T cells, as demonstrated by the significant increment in the dilution of the CTV dye (**Fig. 3j and Suppl. Fig. 2h**). In summary, our data demonstrate that TIA1 and TIAL1 are needed for T-cell quiescence and, if absent, TIA1 TIAL dKO T cells undergo uncontrolled proliferation, activation and cell death.

### TIA1 and TIAL1 restrain proliferative transcription programs in T cells

TIA1 and TIAL1 shape the cell transcriptome by controlling either the splicing, the abundance and/or the translation of their mRNA targets. Thus, to identify the genes and molecular mechanisms regulated by TIA1 and TIAL1 in T cells, we combined global transcriptome analyses in naïve, effector and memory CD4 and CD8 T cells (**Suppl. Fig. 3a and 3b**) with a deep characterization of the RNA interactome of TIA1 and TIAL1 in naïve and/or *in-vitro* activated T cells using individual crosslinking immunoprecipitation (iCLIP) and sequencing (**Suppl. Fig. 3c and Suppl. Table 2**).

In agreement with the important role of TIA1 and TIAL1 in the regulation of RNA splicing, stability and translation, we found TIA1 and TIAL1 mostly associated with U-rich elements present in introns and 3’ untranslated regions (3’UTRs) of both naïve and activated T cells (**Suppl. Fig. 3d, 3e and 3f**). TIA1 and TIAL1 binding density was higher in 3’UTRs, allowing detection of more protein crosslinks sites and targeted genes with high confidence (**Suppl. Fig. 3f, 3g and 3h**). TIAL1 iCLIP assays in naïve T cells yielded a poorer detection of high confident crosslink sites, defined as being annotated by 5 or more unique cDNAs and having an FDR<0.01 (**Suppl. Fig. 3f**). This could be circumscribed to the low mRNA content of naïve T cells compared to activated T cells. Similarly, the low expression of TIA1 limited our iCLIP analysis to activated CD4 T cells and the detection of highly abundant mRNA targets (**Suppl. Fig. 3i**). Despite these difficulties, 1180 genes were annotated as targets of TIA1 in CD4 T cells (**Suppl. Fig. 3j**). Importantly, 1032 of these genes were also targets of TIAL1, reinforcing previous observations on the high binding redundancy of these two proteins ^46, 50^ and allowing us to infer TIA1 binding by just interrogating TIAL1 iCLIP datasets.

In total, we identified 3516 and 7060 genes as targets of TIAL1 in naïve and activated CD4 T cells, respectively (**Suppl. Fig. 3k**). 4053 genes were annotated as TIAL1 targets in naïve CD8 T cells (**Suppl. Fig. 3l**). Identification of TIAL1 targets was largely reproducible, with an up to 87% overlap between naïve and activated CD4 T cells when targeting occurred in 3’UTRs (**Suppl. Fig. 3k**). Importantly, 817 out of the 1156 genes for which TIAL1 crosslinks were found in 3’UTRs in naïve CD8 T cells were also detected as targets in iCLIP assays performed in activated CD4 T cells. This was in the range of variability observed between independent TIAL1 iCLIP experiments, in which the overlap in detected target genes ranged from 70 to 90% depending on the stringency of detection (**Suppl. Fig. 3m**), and prompted us to perform downstream analyses with the richest TIAL1 iCLIP data collected from activated CD4 T cells. This was reinforced by the fact that more than 75% of genes detected as TIAL1 targets in *in-vitro* activated T cells were expressed in all peripheral T cell subsets. Importantly, this percentage increased to almost 100% when TIAL1 crosslink sites were annotated to the 3’UTR (**Suppl. Fig. 3n**). Altogether, we conclude that TIAL1 iCLIP assays performed in *in-vitro* activated CD4 T cells largely recapitulates the protein:RNA interactions occurring in naïve and antigen-experienced peripheral T cells.

Genes associated to the spliceosome and the ribosome were particularly enriched amongst those identified as targets of TIA1 and TIAL1, underscoring, once again, the importance of these proteins in the cellular regulation of mRNA splicing and translation (**Suppl. Fig. 3o**). Additionally, we identified a significant enrichment of target genes associated to the cell cycle, cell senescence and TCR signalling, independently of the iCLIP data source from naïve or activated T cells. This, along with our comprehensive mapping of the protein:RNA interactome, led us to conclude that TIA1 and TIAL1 might control cell activation and proliferation in T cells by targeting key pathway-specific genes.

To assess the functional consequences of TIA1 and TIAL1 deletion in the transcriptome of T cells, we sorted naïve, effector and memory CD4 and CD8 T cells from the spleen of control and *Tia1^fl/fl^ Tial1^fl/fl^ CD4^Cre^* mice for mRNA isolation and sequencing. We found ample differences in the transcriptome of both CD4 and CD8 T cells in the absence of TIA1 and TIAL1. 1632 and 2195 genes were differentially expressed in naïve and EP CD4 dKO T cells, respectively (**Fig. 4a, Suppl. Table 1**). The transcriptome of CD8 dKO T cells also differed largely when compared to control CD8 T cells. We identified 656, 455, 487 genes to be differentially expressed in naïve, MP and EP CD8 T cells in the absence of TIA1 and TIAL1 (**Fig. 4b, Suppl. Table 1**). Interestingly, we noticed that a majority of differentially expressed (DE) genes were increased in both CD4 and CD8 dKO T cells (**Fig. 4a and 4b**), including those DE genes targeted by TIAL1 (**Fig. 4c**). Further integration of RNAseq and TIAL1:RNA interactome data revealed a global increase in the expression of TIAL1 target genes in naïve dKO T cells (**Suppl. Fig. 4a and Suppl. Table 3**). This could be due to a direct effect of TIA1 and TIAL1 deletion, an indirect effect associated to the overactivation of T cells or both. In summary, our data suggest that TIA1 and TIAL1 limit the overall abundance of their RNA targets in T cells.

**Figure 4.**
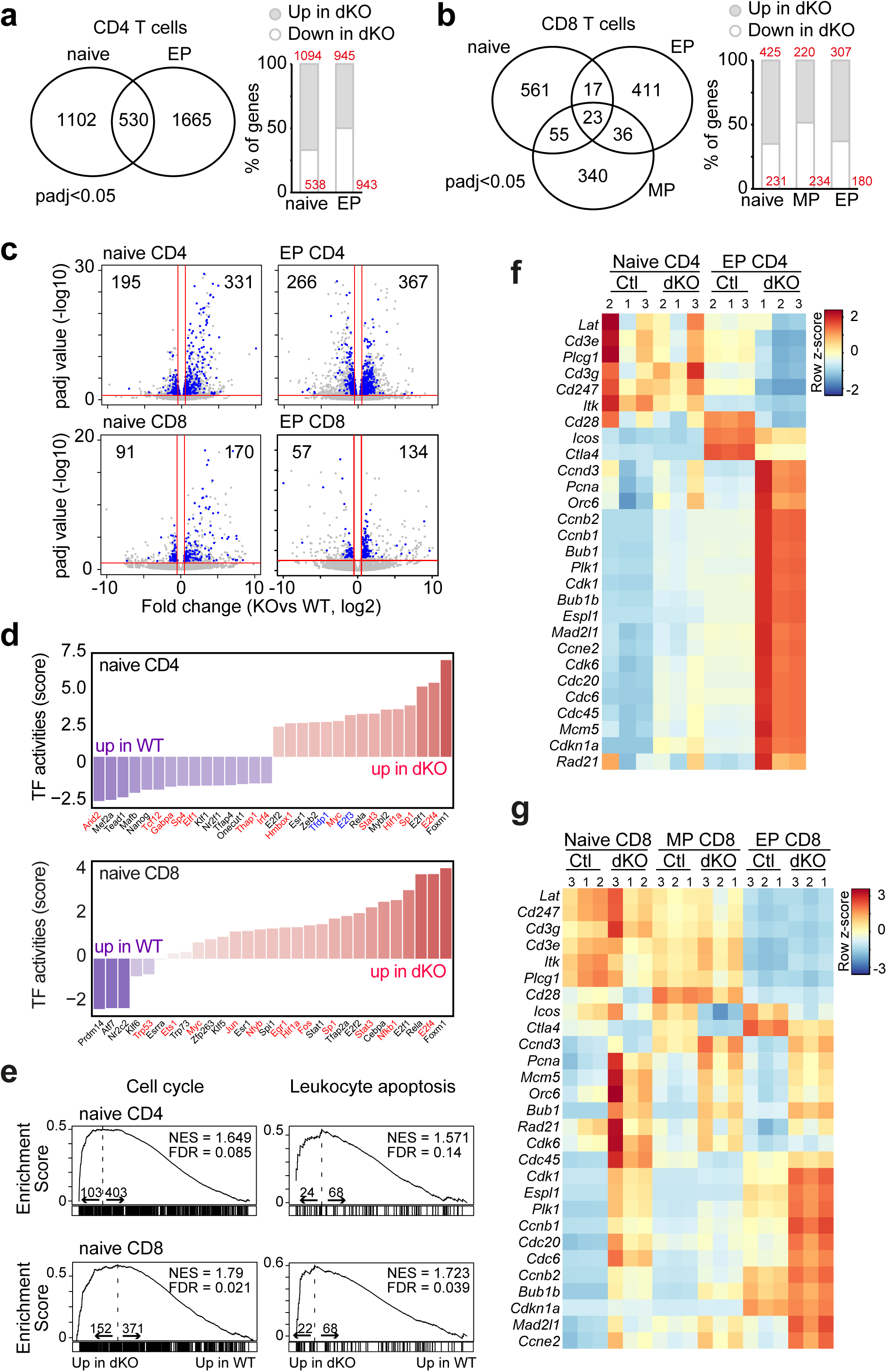
TIA1 and TIAL1 restrain the expression of cell cycle genes. **a,** Number of genes differentially expressed in naïve and EP CD4 T cells in the absence of TIA1 and TIAL1. **b,** Genes differentially expressed in naïve, MP and EP CD8 T cells from *Tia1^fl/fl^ Tial1^fl/fl^ CD4^Cre^*mice. Right panels in a and b show the percentage and number of differentially expressed genes that are up- or down-regulated in TIA1 TIAL1 dKO T cells compared to control T cells. **c,** Volcano plot showing changes in fold expression and p-adjusted (padj) values of genes targeted by TIAL1 and differentially regulated in TIA1 TIAL1 dKO T cells. Up- or downregulated genes are highlighted in blue in each panel. **d,** Inference of TF activities in naïve CD4 and CD8 T cells from control and *Tia1^fl/fl^ Tial1^fl/fl^ CD4^Cre^* mice (performed with decoupleR package). **e,** Gene enrichment plots showing changes in the expression of genes annotated to the cell cycle (GO:0007049) and apoptosis (GO:0071887) in naïve CD4 and CD8 dKO T cells. **f, g,** Heatmaps plot showing the expression of cell cycle and activation genes in CD4 T cells (f) and CD8 T cells (g). RNAseq data were collected from 3 biological replicates per genotype and cell type. P adjusted values (padj) were calculated after Benjamini and Hochberg test correction.

Inference of transcription factor (TF) activities in control and dKO T cells highlighted a global upregulation of genetic programs associated to activation and proliferation of T cells (**Fig. 4d**). Transcription associated to NFkB1/RELA, SP1, STAT3, MYC and E2Fs was significantly increased in naïve dKO T cells. Some of these TFs were identified as targets of TIAL1. However, in almost all cases, their mRNA abundance was unaltered in dKO T cells (**Fig. 4d and Suppl. Table 3**). This suggested that TIA1 and TIAL1 either control the translation of these mRNAs or modulate upstream genetic programs that block their TF activity. Gene set enrichment analyses confirmed the over-representation of pathways associated to the cell cycle and apoptosis in naïve TIA1 TIAL1 dKO T cells (**Fig. 4e and Suppl. Table 4**). Cyclins (*Ccnd3*, *Ccne2*, *Ccnb1* and *Ccnb2*), cyclin-dependent kinases (*Cdk6*) and essential regulators for DNA replication (*Cdc6*, *Cdc20* and *Pcna*) were found amongst the TIAL1-target genes upregulated in dKO T cells compared to control cells (**Fig. 4f**, **Fig. 4g, and Suppl. Table 3**). Taken together, our data demonstrate the direct contribution of TIA1 and TIAL1 on restraining the genetic programs that control T-cell cycling and proliferation in CD4 and CD8 T cells.

### RNA splicing is altered in the absence of TIA1 and TIAL1

TIA1 and TIAL1 are major splicing factors involved in exon definition and inclusion ^45, 50^. Indeed, analysis of the TIAL1:RNA interactome in CD4 T cells showed that these proteins binds, preferentially, U-rich elements found downstream of the 5’ splice site (**Fig. 5a**). So, we asked the question of whether defects in TIA1 TIAL1 dKO T cell homeostasis were due to alteration in mRNA splicing. To test this possibility, we used Multivariate Analysis of Transcript Splicing (rMATS) ^51^ and search for changes in mRNA splicing in control versus TIA1 TIAL1 dKO T cells. Briefly, 1679 and 2022 alternative splicing (AS) events affecting to 1263 and 1559 genes were identified in naïve CD4 and CD8 TIA1 TIAL1 dKO T cells, respectively (**Fig. 5b and 5c, Suppl. Table 5**). Two-thirds of AS events showed a reduction in their inclusion levels of at least 10%, being this an indicator of defective exon definition and inclusion in naïve TIA1 TIAL1 dKO T cells. RNA splicing was also altered in EP TIA1 TIAL1 dKO T cells, but to a lesser extent than in naïve T cells. 343 and 1293 AS events were identified in EP CD4 and CD8 TIA1 TIAL1 dKO T cells, respectively. This was a 5-fold and a 1.5-fold reduction compared to the number of AS events found in naïve CD4 and CD8 TIA1 TIAL1 dKO T cells. In addition, we found little overlap (30%) in the number of AS genes identified in naïve and EP T cells in the absence of TIA1 and TIAL1 (**Fig. 5d and 5e**). This could be explained, at least in part, due to the extensive changes in RNA splicing taking place after T-cell activation ^52^.

**Figure 5.**
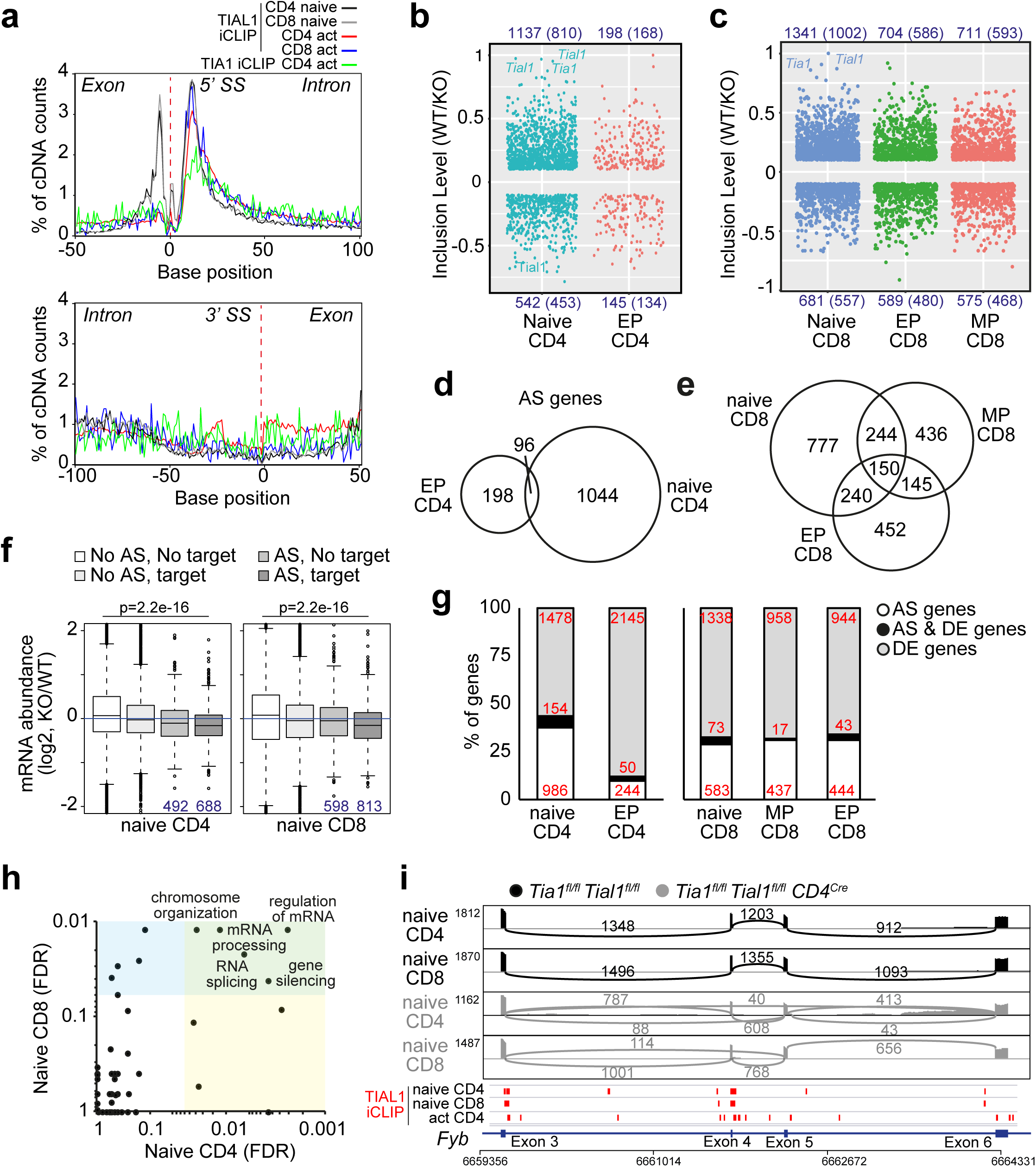
TIA1 and TIAL1 act as splicing regulators in T cells. **a,** RNA map showing the genome location of TIAL1 crosslink sites annotated to exons and introns. **b,** Dot plot of alternative spliced (AS) events detected in naïve and EP dKO CD4 T cells as compared to control cells. **c,** AS events annotated in naïve, MP and EP CD8 T cells in the absence of TIA1 and TIAL1. In b and c, the number in blue indicates the total number of AS with an inclusion level higher or lower than 0.1 (10%) and FDR<0.05. The number in parenthesis indicates the number of genes with altered splicing. **d, e,** Venn diagrams showing the overlap of AS genes in naïve, MP and EP T cells in the absence of TIA1 and TIAL1. **f,** Changes in mRNA abundance of TIAL1 target genes undergoing alternative splicing in naïve dKO T cells. In blue, number of AS genes in each group. Kolmogorov–Smirnov test was performed for statistical analysis of data distribution. Box plot shows the median, lower and upper hinges corresponding to the first and third quartiles and a 95% confidence interval. **g,** Percentage of AS genes that are also differentially expressed in TIA1 TIAL1 dKO T cells. **h,** Identification of the main biological processes affected by alternative splicing in naïve TIA1 TIAL1 dKO T cells. Pathway enrichment analyses were performed with Webgestalt. **i,** Sashimi plot showing AS of *Fyb* exon 4 and 5 in naïve CD4 and CD8 T cells in the absence of TIA1 and TIAL1. In red, TIAL1 crosslink sites detected by iCLIP in naive and activated T cells. FDR values were calculated after Benjamini and Hochberg test correction of p values. Values indicate the number of exon-exon spanning reads annotated to each junction.

Globally, changes in splicing in naive TIA1 TIAL1 dKO T cells led to an overall reduction in the RNA abundance of the TIAL1 target genes (**Fig. 5f**). However, at the single gene level, we found little overlap between differentially expressed genes and AS genes (**Fig. 5g**). This suggested that alterations in mRNA splicing in the absence of TIA1 and TIAL1 led to more qualitative than quantitative changes of the transcriptome. Next, we assessed whether changes in RNA splicing could be affecting to genes associated with T-cell homeostasis and/or activation. Despite our search, pathway enrichment analyses highlighted instead several RNA-related biological processes as being most affected by the loss of TIA1 and TIAL1-dependent regulation of mRNA splicing (**Fig. 5h**). We identified some AS events affecting to TCR-signal transducers (e.g. *Fyb*) and chromatin re-modelers (e.g. *Kmt2c*) in both CD4 and CD8 TIA1 TIAL1 dKO T cells (**Fig. 5i and Suppl. Table 5**). Interestingly, we could link certain alternative exon usage occurring only in naïve CD4, CD8 T or in both cell types with the annotation of selective TIAL1 binding in the downstream intron of AS exon. This was exemplified by the differential splicing of the FYN binding protein (*Fyb*) and the Vacuolar Protein Sorting 13a (*Vsp13a*), two proteins involved, respectively, in TCR signalling and T-cell cytolysis in inflamed tissues ^53^. AS of *Fyb* exon 4 was common to naïve CD4 and CD8 TIA1 TIAL1 dKO T cells, and this correlated with the annotation of TIAL1 crosslink sites close to the 5’ splice site of the downstream intron (**Fig. 5i**). By contrast, TIAL1 binding and alternative *Fyb* exon 5 skipping was only detected in naïve CD4 T cells (**Fig. 5i**). Similarly, *Vsp13a* exon 16 was skipped only in naïve CD4 dKO T cells (**Suppl. Fig. 4b**) whereas *Vsp13a* exon 69 was skipped uniquely in naïve CD8 dKO T cells (**Suppl. Fig. 4c**). In both cases, TIAL1 crosslink sites were detected in close proximity to the 5’ splice site of the downstream intron, suggesting that TIA1 and TIAL1 binding and interaction with other splicing regulators might define cell specificity and extension of AS events. In summary, our data highlight that TIA1 and TIAL1 shape qualitatively the transcriptome of T-cells by controlling both common and T-cell subset specific splicing programs, some of which might have an impact on TCR signal transduction and cell activation.

### TIA1 and TIAL1 control the expression of quiescence-associated transcription factors

TIA1 and TIAL1 also control the stability and translation of selected mRNA targets by binding to their 3’UTRs ^41, 43, 46^. In both naïve and *in-vitro* activated T cells, TIA1 and TIAL1 crosslinks were preferentially annotated within the last 50 nucleotides prior to the polyadenylation sequence (**Fig. 6a**). We identified a total of 4094 genes targeted by TIAL1 in 3’UTRs (**Suppl. Fig. 3g**). Amongst these targets, we found 62 TFs that were differentially expressed in naïve and/or EP dKO CD4 T cells (**Suppl. Fig. 5a and Suppl. Table 3**). We mapped TIA1 and TIAL1 crosslink sites along the 3’UTR of *FoxP1, Lef1, Tcf7* and many others TFs controlling T-cell quiescence, maintenance and activation (**Fig. 6b**). The abundance of *FoxP1, Lef1, Egr2, Ikzf1* and *Zeb1* mRNAs was remarkably diminished between 1.5 and 2-fold in naïve dKO CD4 T cells compared to control T cells (**Fig. 6c**). The expression of these quiescence-associated TFs was also decreased in EP dKO CD4 T cells, as well as other important repressors of T-cell activation and differentiation such as *Bach2, Id3, Tcf7* and *Egr2* (**Fig. 6c and Suppl. Table 3**). In deep contrast, several activators of the cell cycle and proliferation, such as *Myb*, *Batf, Bhlhe40, Runx3* and *Eomes,* were found increased in TIA1 TIAL1 dKO CD4 T cells (**Suppl. Fig. 5a and 5b**). This was independent of TIA1 and TIAL1 binding to their transcripts (**Suppl. Fig. 5a and 5b**) and reflected, most likely, the loss of transcriptional repression in the absence of these proteins. Importantly, the reduction in quiescence-associated TFs (e.g. *Foxp1*, *Bach2 and Id3*) and the concomitant increased in activation-associated TFs (e.g. *Irf5, Irf8, Id2*, *Bhlhe40*, *E2F8, Gata1*…) was recapitulated in TIA1 TIAL1 dKO CD8 T cells (**Fig. 6d** and **Suppl. Fig. 6a and 6b**). Altogether, our data highlight that TIA1 and TIAL1 contribute to the establishment of the T-cell quiescence program by securing the expression of TFs repressing T-cell activation and proliferation.

**Figure 6.**
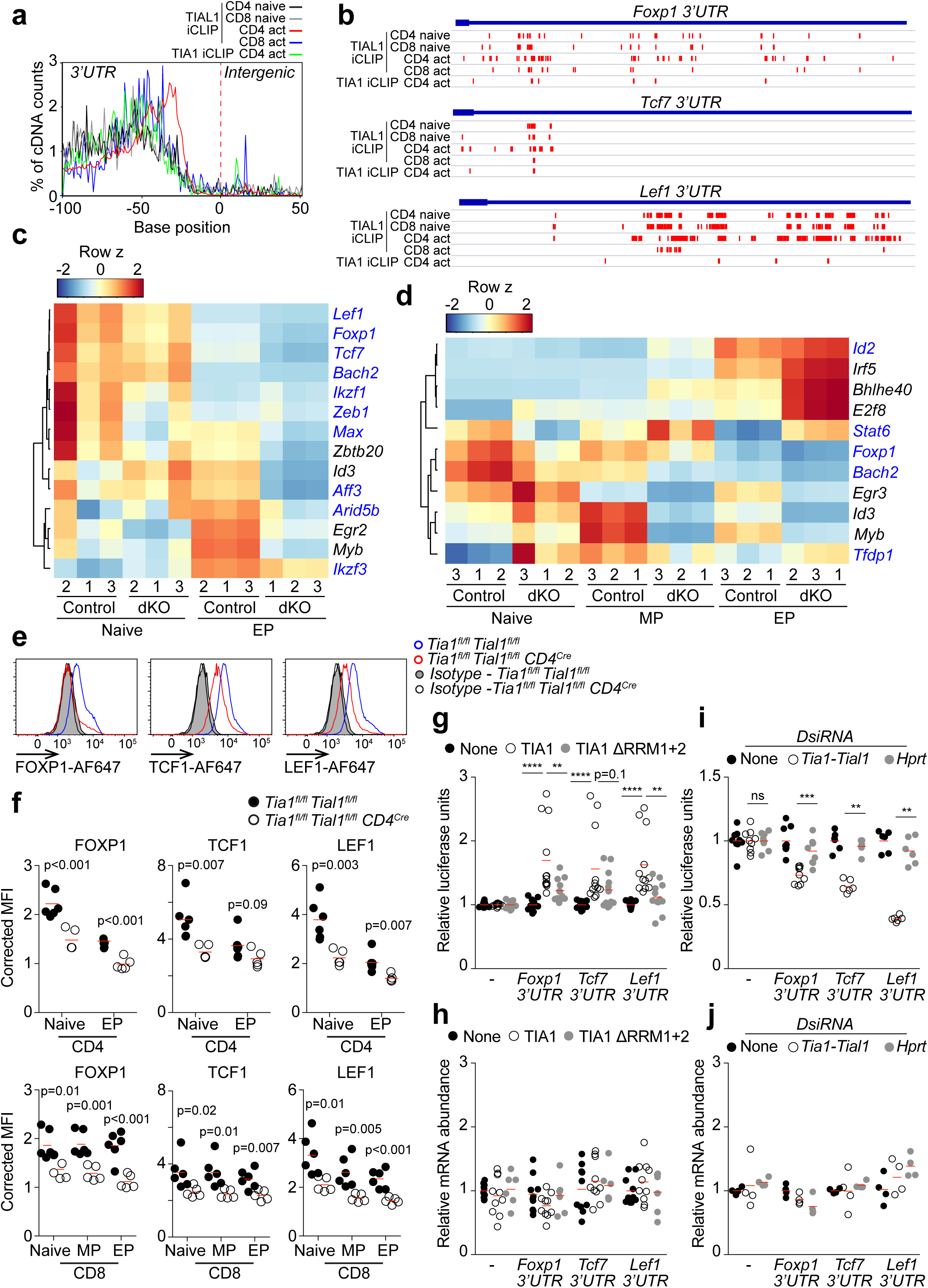
TIA1 and TIAL1 control the transcriptional program for T-cell quiescence. **a,** RNA map showing the genome location of TIA1 and TIAL1 crosslink sites annotated to 3’UTRs in naïve and activated T cells. **b,** Genomic view of TIA1 and TIAL1 crosslink sites annotated within the 3’UTRs of *Foxp1, Tcf7* and *Lef1*. Data from iCLIP experiments performed in naïve and activated T cells are shown. **c,** Heatmap showing the mRNA abundance in CD4 T cells of selected, differentially expressed TFs bound by TIAL1 within their 3’UTR. In blue, TIAL1 targets annotated in both naïve CD4 and CD8 T cells, and in *in-vitro* activated CD4 T cells. In black, TIAL1 targets identified mainly in activated CD4 T cells. **d,** Heatmap showing the mRNA expression in CD8 T cells of TFs targeted by TIAL1 (selected as in c). **e,** Flow cytometry analysis of FOXP1, TCF1 and LEF1 expression in naive CD4 T cells from control and *Tia1^fl/fl^ Tial1^fl/fl^ CD4^Cre^* mice. **f,** Quantitation of FOXP1, TCF1 and LEF1 expression in splenic CD4 and CD8 T cells. MFI was corrected by the MFI of an isotype antibody control. Data are representative from one of the three independent experiments performed with n=5 mice per genotype. **g,** Comparative analysis of the expression of renilla luciferase reporters under the control of the 3’UTR of *Foxp1*, *Tcf7* and *Lef1* after co-transfection with a wild-type TIA1 or a binding-mutant TIA1 ΔRRM1+2. **h,** Quantitation by RT-qPCR of renilla luciferase mRNA abundance in transfected HEK 293T cells from g. **i,** Analysis of renilla luciferase reporter expression in HEK293T cells after knock down of TIA1 and TIAL1 using DsiRNAs. **j,** Quantitation of renilla luciferase mRNA abundance in TIA1-TIAL1 knock down HEK293T cells shown in i. No-DsiRNA treated or Hprt-knock down cells were used as negative control. In g and I, relative luciferase units were calculated as renilla luciferase units normalized by firefly luciferase units to account for changes in cell transfection efficiency. In h and j, data was further normalized by the expression of an 18S rRNA to account for initial changes in RNA concentration and retro-transcription into cDNA. Data is pooled from two (i, j) or three (g,h) independent experiments performed by duplicate or triplicate. Mann-Whitney tests are performed for statistical analysis of the data in f to j.

Assessment by flow cytometry of FOXP1, TCF1 and LEF1, three essential TFs controlling T-cell quiescence, confirmed the reduction of these proteins in naïve, MP and EP CD4 and CD8 T cells from *Tia1^fl/fl^ Tial1^fl/fl^ CD4^Cre^* mice (**Fig. 6e** and **6f**). To gather mechanistic insight into how binding of TIA1 and TIAL1 controlled the expression of FOXP1, TCF1 and LEF1, we generated reporter assays in which each of the 3’UTR of *FoxP1, Tcf7* and *Lef1* were cloned downstream the coding sequencing of a renilla luciferase reporter. Overexpression of a wild-type isoform of TIA1, but not of a mutant TIA1 lacking the RNA recognition motifs 1 and 2, promoted mRNA translation of each of these reporters in transiently transfected HEK293 cells without altering mRNA abundance (**Fig. 6g** and **6h**). Complementary, knock down of both TIA1 and TIAL1 significantly decreased the mRNA translation, but not the mRNA levels, of all the luciferase reporters under the regulation of the 3’UTR of FoxP1, Tcf7 or Lef1 to a different degree (**Fig. 6i** and **6j, Suppl. Fig. 6c**). Assessment of the mRNA stability of the renilla luciferase reporters normalised to a firefly luciferase mRNA internal control showed no changes in mRNA abundance after TIA1 overexpression (**Suppl. Fig. 6d)**. In summary, our data demonstrate that TIA1 and TIAL1 binding to the 3’UTR of *FoxP1, Tcf7 and Lef1* mRNAs preferentially control their translation into protein.

### FOXP1 quiescent program is impaired in the absence of TIA1 and TIAL1

In naïve T cells, FOXP1 is a primary regulator of quiescence that acts as a repressor of downstream TFs modulating T-cell activation and proliferation ^54, 55^. Thus, we first confirmed by western blotting the reduction in FOXP1 expression in naïve CD4 T cells from *Tia1^fl/fl^ Tial1^fl/fl^ CD4^Cre^* mice observed by flow cytometry (**Fig. 7a**). Then, to assess if the FOXP1 transcriptional program was altered in TIA1 TIAL1 dKO T cells, we reanalysed a previously published FOXP1 ChIPseq dataset ^55^ and identified the FOXP1 target genes induced or repressed in naïve T cells when compared to EP T cells (**Suppl. Fig. 7a and Suppl. Table 6**). Global analysis of these FOXP1 target genes in naïve CD4 and CD8 TIA1 TIAL1 dKO cells revealed a general increase in the expression of genes repressed by FOXP1 whereas those genes induced by FOXP1 were significantly reduced when compared to control naïve T cells (**Fig. 7b**). These changes were independent of TIAL1 binding to the mRNA transcripts of genes regulated by FOXP1 (**Fig. 7c**), suggesting that direct binding and regulation of FOXP1 by TIA1 and TIAL1 is the main driver of the transcriptional remodelling occurring in naïve TIA1 TIAL1 dKO cells.

**Figure 7.**
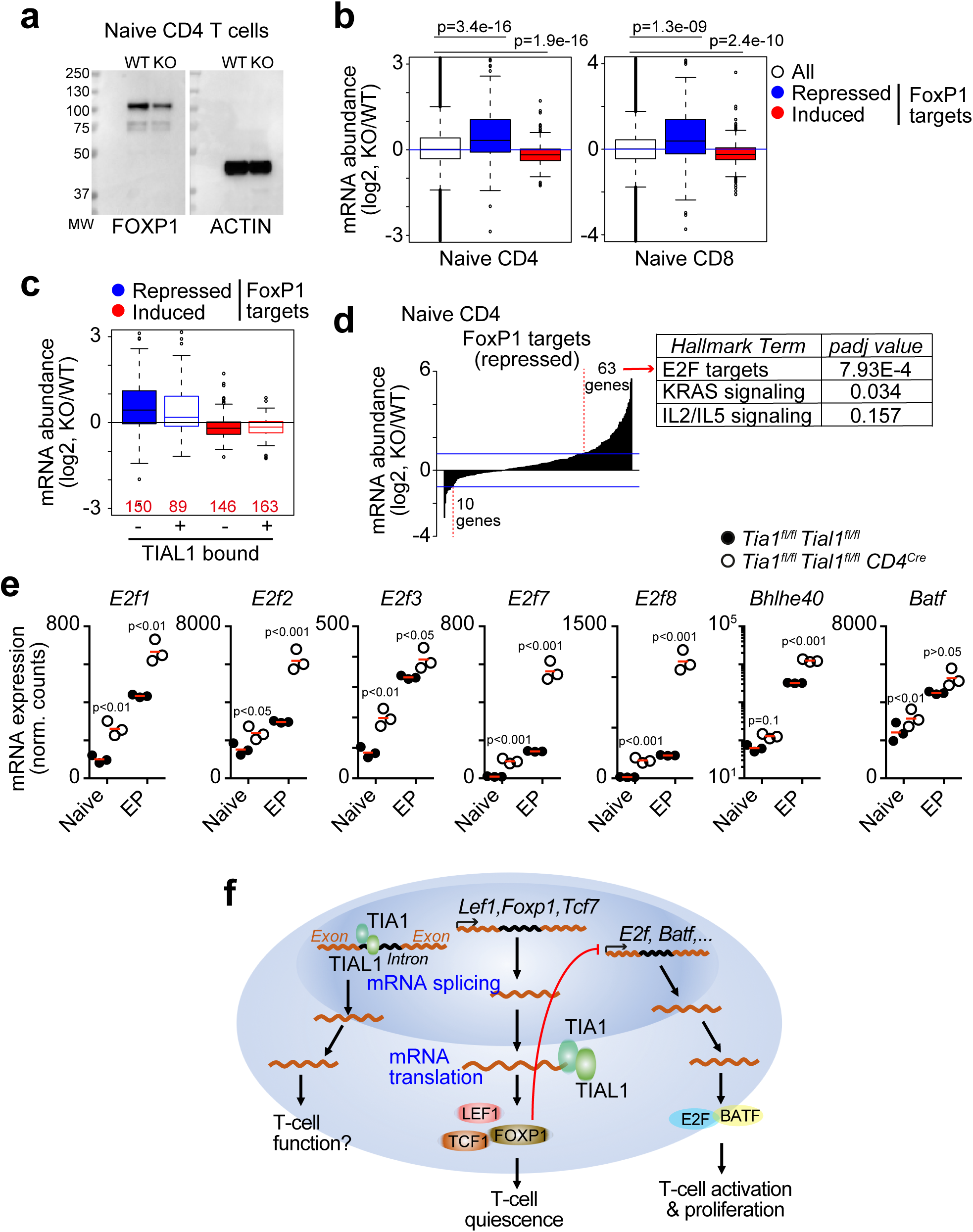
The quiescence program triggered by FoxP1 is deregulated in TIA1 TIAL1 dKO T cells. **a,** Immunoblot detecting FOXP1 and ACTIN in control and dKO naïve CD4 T cells. **b,** Global expression changes of genes transcriptionally regulated by FOXP1 in naïve CD4 and CD8 T cells. ChIPseq data (GSE121279) was analysed to identify FOXP1 target genes induced or repressed in naïve T cells when compared to EP T cells. Wilcox tests were performed in the statistical analyses. **c,** Comparison of the changes in expression of FOXP1 target genes which transcripts are bound, or not, by TIAL1 in naive CD4 T cells. In red, the number of genes in each group. Box plots in b and c show the median, lower and upper hinges corresponding to the first and third quartiles and a 95% confidence interval. **d,** Bar plot showing the number of FOXP1-target genes with changes in expression higher than 2-fold in naïve CD4 dKO T cells. Right panel, pathways enriched in highly upregulated FOXP1 target genes in naïve CD4 dKO T cells. **e,** RNAseq expression analysis of the indicated genes in naïve and EP CD4 T cells from control and *Tia1^fl/fl^ Tial1^fl/fl^ CD4^Cre^* mice. Mann-Whitney tests were performed for statistical analysis. **f,** Proposed model by which TIA1 and TIAL1 contribute to the maintenance of peripheral T cells by promoting the quiescence transcriptional program induced by FOXP1.

Among the genes repressed by FOXP1, we identify 63 and 58 genes that were increased by more than 2-fold in naïve CD4 and CD8 cells from *Tia1^fl/fl^ Tial1^fl/fl^ CD4^Cre^* mice respectively (**Fig. 7d and Suppl. Fig. 7b**). Pathway enrichment analyses showed that these genes were associated with the KRAS and E2F pathways. FOXP1 is a major repressor of E2F TFs and blocks cell cycle entry in the absence of antigen encounter ^6^. Assessment of E2F family members showed a significant increase in the expression of *E2f1, E2f2, E2f3, E2f7, E2f8* genes in TIA1 TIAL1 dKO T cells (**Fig. 7e and Suppl. Fig. 7c**). Expression of *Bhlhe40* and *Batf*, two activation-associated TFs repressed by FOXP1, was also increased in naïve and/or EP T cells from *Tia1^fl/fl^ Tial1^fl/fl^ CD4^Cre^*mice. We identified most of *E2f*-family members, *Bhlhe40* and *Batf* mRNAs as targets of TIAL1 when expressed abundantly in activated T cells (**Suppl. Fig. 7d and Suppl. Table 2**). However, luciferase reporter assays revealed that binding of TIA1 and TIAL1 to the 3’UTR of *E2f3, Bhlhe40* and *Batf* was largely dispensable for their expression (**Suppl. Fig. 7e**). Taken together, TIA1 and TIAL1 act as selective post-transcriptional regulators of FOXP1 expression and, if absence, the FOXP1-associated quiescence program is impaired leading to anomalous T-cell activation, proliferation and cell death.

## DISCUSSION

Mounting evidences highlight the crucial roles of post-transcriptional regulation by RBPs on shaping T-cell helper and cytotoxic responses ^20, 26^. Here, we show that RBPs are also essential for the maintenance of naïve T cells by enforcing quiescence. We demonstrate that TIA1 and, its orthologous, TIAL1 are dispensable for late T cell development in the thymus, but most needed for the maintenance of CD4 and CD8 T cells in peripheral lymphoid organs. TIA1 and TIAL1 act redundantly in T cells as shown by the presence of a similar number of T cells in single conditional KO mice and control mice. By contrast, deletion of both TIA1 and TIAL1 leads to a high drop in the number of peripheral CD4 and CD8 T cells associated with aberrant activation, proliferation and death in the absence of stimulation with cognate antigens.

Naïve T cells are characterized for having low cellular mass, low energy production and low proliferative capacity. However, naïve T cells are still metabolically active. This is exemplify by the rapid protein turnover of quiescence-promoting TFs, the translational silencing of early activation and cyclin genes (e.g. *Cd69*, *Jun*,…), and the accumulation of non-engaged ribosomal proteins and glycolytic enzymes that fuel T cell growth, proliferation and differentiation upon TCR activation ^11^. Entangled regulation at the transcriptional and post-transcriptional level maintains this state of cell preparedness. Here we demonstrate that TIA1 and TIAL1 control selectively the expression of TFs enforcing quiescence and, in their absence, T cells proliferate and die, possibly, due to exhaustion. Mitochondrial energy metabolism and overall mRNA translation are comparable in peripheral control and TIA1 TIAL1 dKO CD4 and CD8 T cells. Thus, it is possible that anomalous hyperactivation and proliferation is not metabolically supported in the absence of TIA1 and TIAL1 and this leads to cell death.

Amongst others, we have identified FOXP1, TCF1 and LEF1 as being key quiescence-associated TFs targeted and regulated by TIA1 and TIAL1. TIA1 TIAL1 dKO T cells express very low levels of these TFs, and their phenotype largely recapitulates the one observed in T cells lacking FOXP1, TCF1 or LEF1 ^5, 54, 56, 57, 58, 59^. Our data support previous evidence suggesting that the expression of FOXP1, TCF1 and LEF1 is heavily regulated at the post-transcriptional level ^11, 12^. TIA1 and TIAL1 contribute to the cellular maintenance of *Foxp1*, *Tcf7* and *Lef1* mRNAs which are characterised for having a short half-life due to polyadenylation shortening mediated by BTG1 and BTG2 ^12^. In addition, the proteasome mediates rapid degradation of FOXP1, TCF1 and LEF1 proteins in naïve T cells, with TIA1 and TIAL1 likely contributing to the constant replenishment of the protein pool by promoting mRNA translation. In an elegant study, Wolf et al. identified FOXP1, TCF1 and LEF1 in the top-5 TFs with the shortest protein half-life in naïve CD4 T cells ^11^. This contrasts with the high stability of activation-associated TFs, such as STAT1 and NFkB1, and suggests that shutdown of *Foxp1*, *Tcf7* and *Lef1* transcription and rapid protein degradation control quick reprograming of quiescence T cells upon activation. Thus, TIA1 and TIAL1 contribute to the long-term identity and maintenance of T cells by securing the expression of quiescence TFs including FOXP1, TCF1 and LEF1.

Our comparative analyses of the transcriptome of T cells highlight the loss of the quiescence program led by FOXP1 in TIA1 TIAL1 dKO T cells. FOXP1 is a master TF that binds the promoter of other important quiescence-associated TFs and induces their expression (e.g. *Lef1, Ikzf1, Zeb1*, *Tfap4*…) ^55, 60, 61^. FOXP1 also acts as a primary repressor of the E2F family of TFs which promote DNA synthesis and cell cycle progression ^6, 62, 63, 64^. We found *Lef1, Ikzf1* and *Zeb1* to be downregulated in TIA1 TIAL1 dKO T cells. By contrast, *E2f* genes were highly expressed and highly active in TIA1 TIAL1 dKO T cells when compared to control T cells. This, along with an increment in the transcriptional activity of TFs associated to T-cell activation and proliferation such as NFkB1, SP-1 and FOXM1, explains the overall increase in the expression of cell cycle-associated genes observed in TIA1 TIAL1 dKO T cells. We have not found any correlation between TIA1-TIAL1 targeting and expression of *E2f-family, Batf or Bhlhe40* genes. Our iCLIP assays annotated, mainly in activated T cells, the interaction of TIAL1 with these and other transcripts encoding many cell cycle proteins (e.g. *Pcna, Ccnb1, Ccnd3, Ccne2,…*). However, TIA1 and TIAL1 protein expression does not always correlate with functionality. It is very likely that fine-tuning post-transcriptional regulation of these genes is eclipsed in naïve TIA1 TIAL1 dKO T cells by the strong transcriptional deregulation associated to the loss of T-cell quiescence. In addition, TIA1 and TIAL1 mechanistically act as important regulators for exon definition and splicing in T cells, as previously reported in B cells and different cell lines ^45, 46, 50^. Changes in splicing were only partially conserved in naïve CD4 and CD8 T cells, indicating that TIA1 and TIAL1 might exert cell-specific functions possibly due to differential partnership with other splicing regulators. Analysis of AS genes showed an enrichment of gene sets associated with RNA biology, but not regulation of activation and proliferation. Thus, TIA1/TIAL1-dependent control of quiescence might be independent of their function as splicing regulators.

We also found that TIA1 and TIAL1 bind extensively to the 3’UTR of mRNA targets to control their expression in a selective manner. We have validated *Foxp1*, *Lef1* and *Tcf7* as mRNA targets which translation is directly regulated by TIA1 and TIAL1. However, for other RNA targets, mechanistic validation is still required at a single gene-specific manner. Identification of direct protein:RNA interactions with iCLIP and/or the presence of U-rich binding motifs cannot be extrapolated into the presence of a functional regulatory node in targets such as *Batf*, *Bhlhe40* and *E2F3*. Indeed, our TIA1/TIAL1 knock down and 3’UTR reporter assays suggest that upregulation of these genes in TIA1 TIAL1 dKO T cells might be mostly driven by the lack of FOXP1-dependent gene suppression. In summary, our data support the existence of a novel post-transcriptional mechanism (**Fig. 7f**) by which TIA1 and TIAL1 control the expression of FOXP1 and other quiescence-associated TFs that block naïve T-cell activation and metabolic reprograming. In the absence of TIA1 and TIAL1, the quiescent transcriptional program is impaired enabling the expression of activation-associated TFs (E2Fs, BATF,…) and cell cycle genes for aberrant naïve T cell proliferation and differentiation into a effector-like phenotype.

Recent characterization of the RBPome of human and mouse helper T cells has identified more than 1000 proteins with the capacity to bind RNA ^65^. However, only a handful of RBPs have been implicated in the regulation of T-cell homeostasis. For example, mutation of the RBP Roquin leads to a loss in T-cell homeostasis and autoimmunity ^66^. Roquin-1/2 are highly expressed in naïve and resting memory T cells and repress costimulatory receptors to enforce quiescence (e.g. ICOS and OX40) ^67, 68^. The mechanism by which Roquin-1/2 contribute to naïve T cell maintenance is likely independent of TIA1 ^69^, pointing towards the existence of multiple layers of post-transcriptional regulation in naïve T cells. Indeed, the splicing factor SRSF1 controls T-cell homeostasis by preserving the expression of PTEN that blocks mTOR-dependent translation and activation of naïve T cells ^70, 71^. Similarly, the translational regulator PDCD4 keeps the low metabolic profile of quiescent T cells but, in this case, by targeting the eIF4 complex and blocking protein synthesis. These examples, along with our description of TIA1 and TIAL1 being necessary for the expression of the TFs enforcing quiescence, highlight the co-existence of several post-transcriptional mechanisms that support long-term maintenance and readiness of naïve T cells. In the future, description of new relevant RBPs and the interplay between these post-transcriptional mechanisms in T-cell homeostasis and activation will build the foundations for enhancing novel RNA-based immunotherapies.

## Acknowledgments

We thank to all personnel from the animal facility of Toulouse - CREFRE - and from the flow cytometry (F. L’Faqihi, V. Duplan-Eche and AL. Iscache), transcriptomics (A. Chaubet) and bioinformatics technical platforms of INFINITy. This project was supported by the ATIP-Avenir, Plan Cancer program (C18003BS), the French National Research Agency, ANR (ANR-20-CE15-0007 and ANR-22-CE15-0013), FRM Foundation (EQU202303016269), La Ligue contre le cancer (R21011BB) and Cancéropôle Grand Sud-Ouest (R20067BB).

## CRediT author contributions

Ines C. Osma-Garcia: Conceptualisation, Investigation, Formal analysis, Writing - review and editing. Orlane Maloudi: Conceptualisation, Investigation, Formal analysis, Writing - review and editing. Mailys Mouysset: Conceptualisation, Investigation, Formal analysis, Writing - review and editing. Dunja Capitan-Sobrino: Investigation, Formal analysis. Trang-My M. Nguyen: Investigation, Formal analysis. Yann Aubert: Formal analysis, Writing - review and editing. Manuel D. Diaz-Muñoz: Conceptualisation, Investigation, Formal analysis, Resources, Supervision, Funding acquisition, Writing — original draft, review and editing, Project administration. As first co-authors in this manuscript, Ines C. Osma-Garcia, Orlane MaloudI and Mailys Mouysset might swap their names as convenient for career development purposes.

## Ethics declaration

The authors declare no competing interests.

## METHODS

### Mice

*Tia1^tm1c(KOMP)Wtsi^* (*Tia1^fl/fl^*) and *Tial1^tm1c(EUCOMM)Wtsi^* (*Tial1^fl/fl^*) mice were generated as previously indicated ^45^ and crossed with transgenic Tg(Cd4-cre)^1Cwi/BfluJ^ for conditional gene deletion in T cells. Mice were maintained on a C57BL/6 background. Randomization, but not experimental ‘blinding’, was set in these studies by housing *Tia1^fl/fl^ Tial1^fl/fl^ CD4^Cre^*mice and Cre-negative littermate control mice in the same cage from weaning. Both females and males of 8–16 weeks of age were used in all experiments. No primary pathogens or additional agents listed in the FELASA recommendations were confirmed during health monitoring surveys of the mouse stock. Ambient temperature was ∼19-21C and relative humidity 52%. Lighting was provided on a 12-hour light: 12-hour dark cycle. After weaning, mice were transferred to individually ventilated cages with 2-5 mice per cage. Mice were fed ad libitum and received environmental enrichment. All experimental procedures were approved by the local ethical committee of INFINITy and by the French Ministry of Education, Research and Innovation.

### T cell isolation and cell culture

Naïve T cells were isolated from the spleen and/or lymph nodes (axillar, brachial and inguinal LNs) by smashing these tissues in 40 μm cell strainers containing 5-10 ml of complete RPMI-1640 medium (Dutch Modification from Thermo Scientific supplemented with 10% FCS, 100 U/mL penicillin, 10 μg/ml streptomycin, 2 mM L-glutamine, 1 mM sodium pyruvate and 50 μM β-mercaptoethanol) with the help of a needle plunger. ACK Lysis buffer (Thermo Scientific) was used to lyse red blood cells before T cell isolation using the naive CD4^+^ T Cell Isolation Kit from Miltenyi Biotec or the MojoSort™ Mouse CD4 or CD8 Naïve T Cell Isolation Kits from BioLegend. Cells were cultured at a density of 0.5x10^6 cells/ml in complete RPMI medium and stimulated with plate-bound αCD3 (10 μg/ml, clone 145-2C11, BioLegend) and soluble αCD28 (2 μg/ml, clone 37.51, BioLegend) antibodies. *In-vitro* CD4 T cell survival assays were performed in complete RPMI medium supplemented with recombinant murine IL-2 (100 ng/ml, Miltenyi Biotec) or IL-7 (10 ng/ml, Peprotech). Alternatively, splenocytes were cultured at a density of 0.5x10^6 cells/ml in complete RPMI medium and treated with 0.2 or 2 μg/ml of αCD3 or the indicated doses of recombinant murine IL-2, IL-7 and IL-15 (Peprotech).

### Generation of 3’UTR luciferase reporters and cell transfection

3’UTR nucleotide sequences (*Lef1*, NM_010703.5; *Tcf7*, NM_001313981.2; *Foxp1*, NM_053202.2, *Batf,* NM_016767.2*; Bhlhe40, NM_011498.4; and E2F8,* NM_010093.3) were synthesized as GeneBlocks (IDT technologies) and cloned downstream the coding sequence for renilla luciferase of the psiCheck2 dual luciferase reporter plasmid from Promega. MIGR1-empty, MIGR1-TIA1 and MIGR1-TIA1 ι1RRM1+2 plasmids used as control or to overexpress a wild-type or an RNA-binding deficient isoform of TIA1 were described previously ^41^. Plasmids were transfected into HEK293T cells using Lipofectamine 2000 as recommended by the manufacturer (Thermo Fisher Scientific). HEK293T cells were maintained in DMEM media supplemented with 5% FCS, 100 U/mL penicillin, 10 μg/ml streptomycin and 2 mM L-glutamine. For mRNA stability analysis, 5 µg/ml actinomycin D (ActD, Sigma Aldrich) was added to the cell culture medium to block transcription. Renilla and firefly luciferase signal were measured 48h after cell transfection using a Dual-Luciferase® Reporter Assay System from Promega. For TIA1 and TIAL1 knock down in HEK293T cells, we used TriFECTa DsiRNA, Human TIA1 (hs.Ri.TIA1.13) and TriFECTa DsiRNA, Human TIAL1 (hs.Ri.TIAL1.13) kits from IDT Technologies. Negative control (NC) and Hprt DsiRNA are provided with the kits. HEK293T cells were transfected to incorporate the DsiRNA mix using Lipofectamine RNAiMAX as recommended by the manufacturer (Thermo Fisher Scientific). After 48h, DsiRNA and psiCheck2 reporters were co-transfected using Lipofectamine 2000. Total RNA and renilla and firefly luciferase signal were measured after 24h incubation at 37oC. Relative luciferase units were calculated by dividing renilla luciferase counts by firefly luciferase counts to correct for any differences in transfection efficiency. Later, data was relatively quantified to the control group. At least three independent experiments were performed by triplicate. In parallel, RNA was extracted from transfected cells using TriZol (Thermo Fisher Scientific) to measure RNA abundance by RT-qPCR as described previously ^41^. Pooled data from all experiments is shown in each figure panel. Mann-Whitney tests were performed for statistical analyses.

### Flow cytometry

Flow cytometry analyses of T cell subsets in the thymus and peripheral lymphoid organs were performed using the antibodies indicated in **Supplementary Table 7**. Cells were incubated with the Zombie NIR Fixable Viability dye and the Fc Receptor Blocking antibody (clone 2.4G2, BD Biosciences) in PBS+2% FCS (FACs buffer) for 15 min. at 4°C prior cell antigen surface staining. Cells were incubated for 30 min. at 4°C with the selected mix of antibodies in FACs buffer. After extensive washing, cells were fixed and permeabilized for 30 min. at 4°C with the BD Cytofix/Cytoperm™ Fixation and Permeabilization Solution (BD Biosciences) or with the Foxp3 Fix/Perm Buffer Set (BioLegend). Intracellular protein staining was performed in permeabilization buffer containing 1% FCS for at least 1 hour at 4°C. For assessment of cell apoptosis, cells were incubated with CaspACE™ FITC-VAD-FMK in Situ Marker (1 uM per 10^7 cells) for 15 minutes at 37°C in complete RPMI medium prior cell staining with antibodies and Zombie NIR Fixable Viability dye. BrdU labelling of proliferating T cells was assessed after mouse feeding with BrdU (0.8 mg/ml in the drinking water, Sigma Aldrich) for 5 days. Staining for flow cytometry was performed using the BrdU Staining Kit from Thermo Scientific following the instructions from the kit. CellTrace™ Violet Cell Proliferation Kit was used to assess cell proliferation. Cell viability was measured upon staining with the Zombie NIR Fixable Viability dye (BioLegend) prior further staining of surface and intracellular proteins. Sphero AccuCount Beads (ACBP-50-10) were used for counting *in-vitro* cultured cells following instructions from the manufacturer (Spherotech, Inc). Mitochondrial membrane potential was measured using MitoTracker™ Deep Red FM dye and MitoTracker Orange CM-H2TMRo dye (Thermo Fisher Scientific) as previously described ^72^. Briefly, cells were incubated at 37°C in complete RPMI media for 30 min before adding Mitotracker dyes at a final concentration of 1μM. Cells were incubation for another 15 min. prior protein staining for flow cytometry. Measurement of puromycin incorporation into nascent peptides was performed by adding puromycin (10 μg/ml, Sigma Aldrich) with or without harringtonine (2 μg/ml, Sigma Aldrich) to cells previously incubated at 37°C for 30 min. After 7 minutes exactly, cells were placed on ice for surface protein staining, fixation and intracellular staining of puromycin-labelled peptides with a specific antibody coupled to FITC. To evaluate STAT5 phosphorylation, total splenocytes were left in supplemented RPMI medium without FCS at 37°C for 30 min prior adding IL-7 (10 ng/ml). After 30 min. incubation at 37°C, cells were directly fix and permeabilised by adding 1/5 BD™ Phosflow Fix Buffer, 15 min at RT, followed by cell wash, permeabilization with BD™ Phosflow Perm Buffer III, O/N at 4°C and protein staining as described above. Flow cytometry data were collected using a BD Fortessa cytometer and analysed using FlowJo v10.

### FACs cell sorting

T cell subsets from *Tia1^fl/fl^ Tial1^fl/fl^* and *Tia1^fl/fl^ Tial1^fl/fl^ CD4*^Cre^ mice were FACs-sorted for transcriptomics and splicing analyses. Briefly, lymphocyte-M (Cedarlane) was used as recommended to eliminate erythrocytes, dead cells and debris from spleens, and to enrich lymphoid cells that were subsequently stained for 30 min. at 4°C with a mix of antibodies and NIR Fixable Viability dye prepared in PBS + 2% FCS + 1 mM EDTA. T cell sorting was performed in a BD FACS Aria Fusion (BD) gating viability dye negative T cell populations as follow: naïve CD4^+^ T cells, TCRβ^+^ CD4^+^ CD8^-^ CD44^-^ CD62L^+^ CD25^-^ NK1.1^-^ GITR^-^, EP CD4+ T cells, TCRβ^+^ CD4^+^ CD8^-^ CD25^-^ CD44^+^ CD62L^-^ NK1.1^-^ GITR^-^; naïve CD8+ T cells, TCRβ^+^ CD4^-^ CD8^+^ CD44^-^ CD62L^+^ CD25^-^ NK1.1^-^ GITR^-^; MP CD8+ T cells, TCRβ^+^ CD4^-^ CD8^+^ CD44^+^ CD62L^+^ CD25^-^ NK1.1^-^ GITR^-^; and EP CD8+ T cells, TCRβ^+^ CD4^-^ CD8^+^ CD44^+^ CD62L^-^ CD25^-^ NK1.1^-^ GITR^-^.

### RNA sequencing library preparation

Total RNA from FACs-sorted T cells was isolated using the RNeasy Micro Kit from Qiagen. RNA quality was analysed on the Bioanalyzer 2100 (Agilent) and quantified in a Qubit 4 Fluorometer (ThermoFisher Scientific). 10 ng. of RNA were then used to prepare RNAseq libraries using the NEBNext® Single Cell/Low Input RNA Library Prep Kit and NEBNext® Multiplex Oligos for Illumina® (Dual Index Primers Set 1) from New England Biolabs. Three RNAseq libraries were generated per genotype and cell type using unique indexes for sample multiplexing. 100 bp paired-end sequencing was performed using the DNBseq platform from BGI Genomics.

### Individual cross-linking immunoprecipitation (iCLIP)

Individual nucleotide cross-linking immunoprecipitation (iCLIP) was performed to annotate the TIAL1:RNA interactome as previously described ^45, 46, 50^. Briefly, naïve CD4 and CD8 T cells were isolated from the spleen and peripheral LNs of C57BL/6 mice using the MojoSort™ Mouse CD4 or CD8 Naïve T Cell Isolation Kit from BioLegend and irradiated with 254 nm-UV light (600 mJ/cm2 using a Stratalinker 2400). Alternatively, naïve T cells were stimulated *in-vitro* with 10 μg/ml plate-bound αCD3 and 2 μg/ml of soluble αCD28. After 48 hours, cells were washed with ice-cold PBS and irradiated as indicated above. Ice-cold RIPA buffer was used for cell lysis during 15 min. at 4°C followed by sonication (10s, x3) for cell lysate clarification. After centrifugation at 15000 rpm, gDNA was removed by treating the cell lysate supernatant with TurboDNAse (10 U/ml of lysate, ThermoFisher Scientific). RNA was partially digested by treating samples with RNase I (0.167 U/ml, ThermoFisher Scientific) for 3 minutes at 37°C. Immunoprecipitation of TIAL1-RNA complexes was done using 3 μg. of αTIAL1 antibody (clone EPR11323B, Abcam) previously coupled to 50 μl of protein G dynabeads (ThermoFisher Scientific) for 1 h. at RT. As negative control, we used an isotype rabbit IgG antibody (clone DA1E). Beads and T-cell lysate were mixed and incubated in rotation at 4°C, O/N. After immunoprecipitation, beads were stringently washed with high-salt washing buffer (50 mM Tris-HCl pH 7.4, 1 M NaCl, 1 mM EDTA, 1% NP-40, 0.1% SDS and 0.5% sodium deoxycholate) and with PNK washing buffer (20 mM Tris-HCl pH 7.4, 10 mM MgCl2, 0.2% Tween-20). Then, 3’end dephosphorylation of the RNA was performed using FastAP alkaline phosphatase (ThermoFisher Scientific) and PNK (New England Biolabs). After further washes, one fourth of the sample was ligated to a pre-adenylated infrared labelled L3-IR-App adaptor (/5rApp/AG <u>ATC</u> GGA AGA GCG GTT CAG AAA AAA AAA AAA /iAzideN/AA AAA AAA AAA A/3Bio/ coupled with IRdye-800CW-DBCO (LI-COR)) using T4 RNA ligase I (New England Biolabs) and PNK ^73^. The rest of the sample was ligated to a non-infrared L3-ATT-App DNA Linker (/5rApp/WN ATT AGA TCG GAA GAG CGG TTC AG/3Bio/) for library preparation. After extensive washes, RNA-protein complexes were separated by SDS-PAGE electrophoresis, transferred into a nitrocellulose membrane, visualized in a LI-COR Odyssey system and extracted by cutting and incubating the nitrocellulose fragment in PK buffer (100 mM This-Cl pH 7.5, 100 mM NaCl, 1 mM EDTA and 0.2 % SDS) with 20 U proteinase K (Roche) at 50°C for 60 minutes. RNA was then isolated using phenol/clorophorm extraction and ethanol precipitation. RNA was retrotranscribed into cDNA using the SuperScript IV reverse transcriptase (ThermoFisher Scientific) and the irCLIP_ddRT_primers 14, 15, 19, 20, 44, 45, 46 and 49 which share a common backbone (/5Phos/WWW-barcode-NNNN GAA TAG GAA GAG CGT CGT GAT/iSp18/GGA TCC/iSp18/TAC TGA ACC GC and unique barcode for multiplexing (14, TAACT; 15, TAACT; 19, AATAC; 20, ACGCA; 44, TACAG; 45, TCATA; 46, TCTAC; and 49, TTCTG). After cDNA purification with Agencourt AMPure XP beads (Beckman), cDNA was circularised with CircLigase II (Epicentre), amplified by PCR using Solexa P5/P7 primers and sequenced in a DNBseq platform from BGI Genomics (100bp, single-end sequencing).

### Bioinformatics

iCLIP data was mapped to the mouse genome GRCm39.109 and analysed as previously described using the iMaps pipeline in Flow (https://app.flow.bio/) ^74^. Positionally-enriched k-mer analysis (PEKA) was used for binding motif analysis ^75^.

For transcriptomics analyses, raw sequencing files were processed using nextflow nf-core/RNAseq pipeline (version 3.2). Reads from different sequencing lanes were concatenated using bash cat utility, trimmed with Cutadapt (v3.4) and Trimgalore (v.0.6.7). The quality of resulting reads was assessed with FASTQC-0.11.9 using default parameters. Then, paired-end reads were aligned to the mouse genome (GRCm39-release 103) and quantified with STAR-2.7.5a (with options --runMode alignReads --outSAMtype BAM SortedByCoordinate --quantMode TranscriptomeSAM GeneCounts --twopassMode Basic, with option --sjdbOverhang 100). Alignment files were indexed using SamTools (v. 1.9).

Differential expression analyses were performed with DESeq2 (v.1.28.1) using default parameters. Conditions included genotype and sample preparation day to control for variation in the data due to these parameters. Changes in gene expression with a p adjusted value (padj), calculated using the Benjamini and Hochberg correction, lower than 0.05 were considered significant. Analysis of differential alternative splicing events was performed with rMATS-4.0.2 and Python-2.7.2 (with options -t paired --readLength 100 --libType fr-firstrand) with the same mouse annotation used for STAR index building and read alignment. Differentially spliced events in pairwise comparison were considered significant if the false discovery rate (FDR) was below 0.05 and have an absolute inclusion level higher than 0.1 (this is equal to a 10% isoform change). Gene ontology enrichment analyses were performed using Webgestalt using default settings ^76^. Selected gene ontology gene sets were from AmiGO. Data from our sequencing datasets and gene sets was extracted and plotted in R (v. 4.2.2) using ggplot2 (v3.2.1).

Inference of transcription factor activities were calculated with decoupleR (version 2.5.0) and CollecTRI (Collection of Transcriptional Regulatory Interactions) ^77, 78, 79^, a meta-resource containing 43175 signed TF–gene interactions for 1186 TFs compiled from multiple sources (ExTRI, HTRI, TRRUST, TFActS, IntAct, SIGNOR, CytReg, GEREDB, Pavlidis, DoRothEA A, NTNU curations). These sources include publicly available high-resolution identification of TF binding sites with ChIP-seq, DNase-seq or other methods, *in silico* prediction of TF–gene interactions, inferred interactions by gene expression and TF binding motifs, and manual curation. CollecTRI retains directionally of TF–gene interactions (activating or repressing) to infer overall TF activity when fed with gene expression data from control and KO cells. decoupleR was run as default retaining high and likely-confident TF-target scoring criteria A and B as described ^79^.

FOXP1 ChIPseq data from conventional CD4 T cells (2 biological replicates, GSE121279) were reanalysed using nextflow (v22.12.0-edge) nf-core chipseq pipeline (v2.0.0, doi: zenodo.org/record/7139814). *Foxp1* KO Treg cells were used as control in the absence of data from CD4 T cells. 50bp paired-end sequencing reads were retrieved from GEO using the *prefetch* and *fasterq-dump* utilities from Sratoolkit (v2.11.3). Quality was assessed using FastQC (v0.11.9). Sequencing adapters were trimmed using Trimgalore (v0.6.7) and aligned to the mouse genome GRCm39 v106 using Bwa (0.7.17-r1188). Duplicate reads were marked using Picard MarkDuplicates (v2.27.4). Alignments from independent sequencing runs were merged using Picard MergeSamFiles and deduplicated before filtering using samtools (v1.15.1). Reads unmapped, duplicated, not aligned primarily or mapping to multiple locations were removed with samtools. Reads containing more than 4 mismatches or with an insert size greater than 2kb were removed using bamtools (v2.5.2). FOXP1 peaks were called using Macs2 *callpeak* (v2.2.7.1) using the mappable genome size of GRCm39 (--macs_gsize). 17773 and 4728 peaks were annotated in each of the biological replicates and merged with bedtools (v. 2-2-29.0). HOMER (v4.10.4) was used to call the closest gene transcription start site (TSS) to generate the list of FOXP1 targets.

### Statistics

Statistics were performed in R (v 4.2.2) or using Prism-GraphPad (v. 7) software. Statistical tests are indicated in each figure legend. Briefly, unpaired t-tests or non-parametric Mann-Whitney tests (two-sided) were used for comparisons between two groups (if not stated otherwise). Kolmogorov–Smirnov test was used to assess changes in the empirical distribution function of two samples. Benjamini and Hochberg test was used for multiple testing and false discovery rate calculation. Hierarchical clustering (Ward’s) was used to measure the similarities between biological samples and genes.

### Data availability

mRNAseq and iCLIP data have been deposited in Gene Expression Omnibus (GEO) with accession codes GSE242817 and GSE241828, respectively. ChIPseq datasets are publicly available (GSE121279). Any other data or material are available upon request.

## SUPPLEMENTAL INFORMATION

**Supplementary Figure 1.**
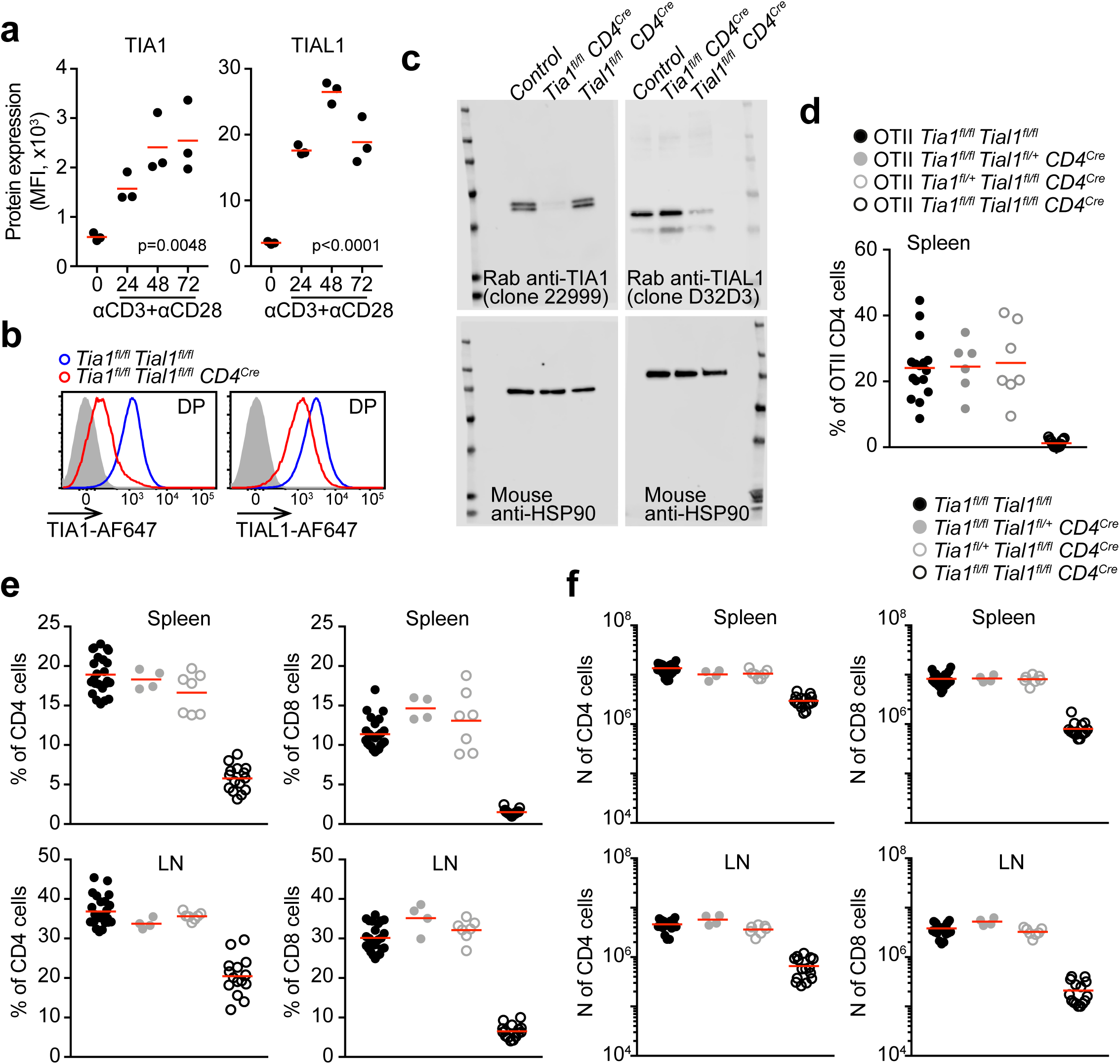
TIA1 and TIAL1 expression and phenotypic characterization of T-cell conditional mice. **a,** Time course analysis by flow cytometry of the expression of TIA1 and TIAL1 in CD4 T cells stimulated with αCD3 and αCD28. 3 biological samples were analysed per time point. One-way ANOVA was performed for statistical analysis of changes in mean expression. **b,** Representative flow cytometry histograms assessing TIA1 and TIAL1 deletion in DP T cells isolated from control and *Tia1^fl/fl^ Tial1^fl/fl^ CD4^Cre^* mice. **c,** Uncropped immunoblots showing efficient TIA1 and TIAL1 deletion in thymic DP T cells (related to Fig. 1c). **d,** Percentage of OTII CD4 T cells found in the spleen of control, single and double TIA1 TIAL1 KO mice. Data pooled from two independent experiments with a minimum of 6 mice analysed per genotype. **e, f,** Proportion (e) and number (f) of CD4 and CD8 T cells found in the spleen and LNs of control, single and double conditional KO mice for *Tia1* and *Tial1*. For control and *Tia1^fl/fl^ Tial1^fl/fl^ CD4^Cre^* mice, data are from 3 independent experiments with at least 5 mice per group. For single conditional KO mice, data from at least 4 mice per genotype.

**Supplementary Figure 2.**
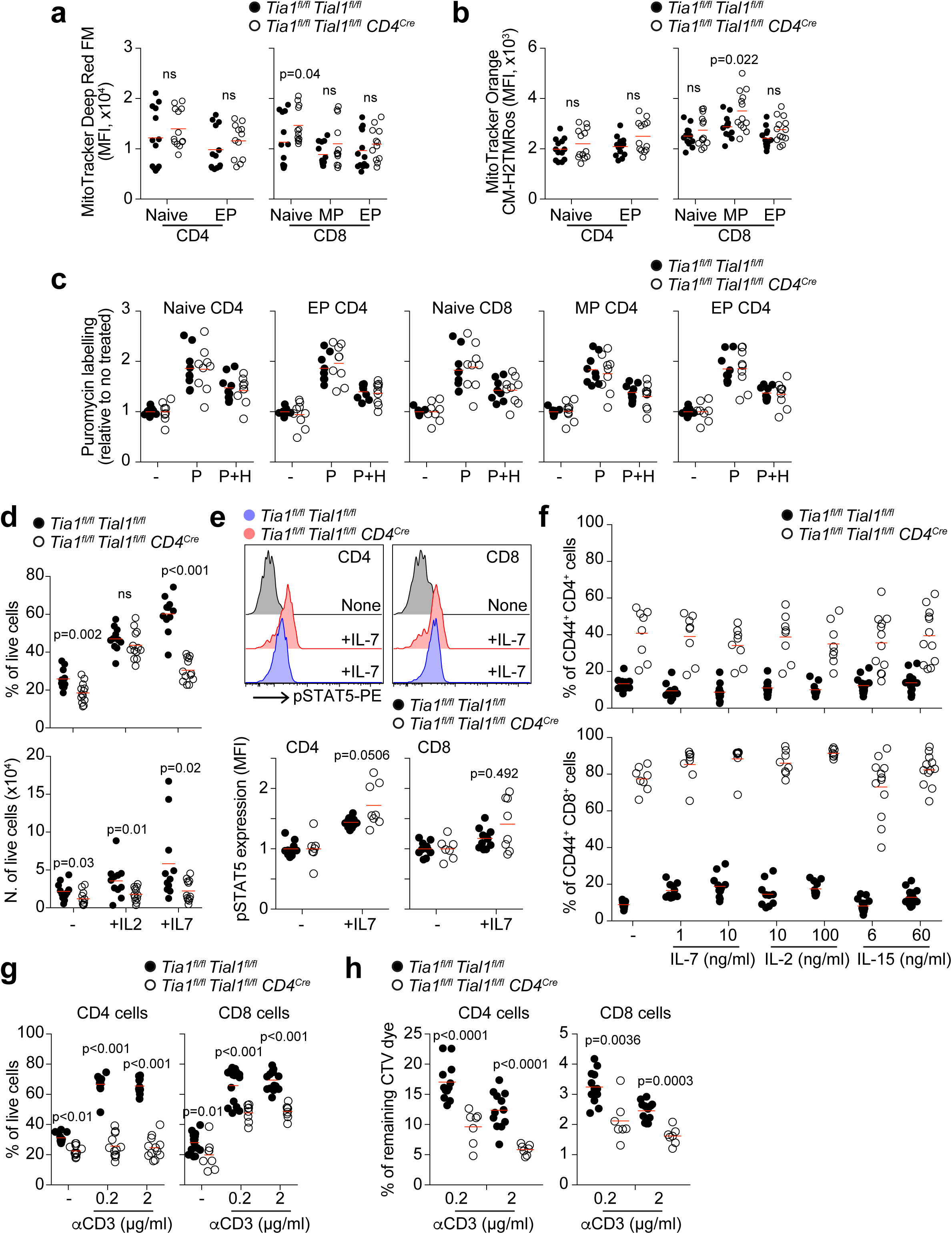
Increased T cell activation and death in the absence of TIA1 and TIAL1. a, b, Analysis of total mitochondrial membrane potential (a) and reducing capacity of mitochondrial membrane potential (b) in *ex-vivo* CD4 and CD8 T cells from control and *Tia1^fl/fl^ Tial1^fl/fl^ CD4^Cre^* mice using the MitoTracker™ Deep Red FM dye and MitoTracker Orange CM-H2TMRo dye. **c,** Quantitation of puromycin (P) incorporation of nascent peptides during mRNA translation in control and TIA1 TIAL1 dKO CD4 and CD8 T cells. The translation initiation inhibitor harringtonine (H) was added to measure peptide synthesis from actively translating ribosomes. **d,** Analysis of naïve CD4 T cell survival after *in-vitro* culture for 24 h. in the absence or the presence of IL-2 or IL-7. **e,** Representative histogram showing pSTAT5 detection in naïve CD4 and CD8 T cells from control and *Tia1^fl/fl^ Tial1^fl/fl^ CD4^Cre^* mice after stimulation with IL-7 at 37°C for 30 min. Bottom panels, quantitation of pSTAT5 relative to the expression in control T cells. **f,** Proportion of CD44^+^ viable T cells after treatment with IL-7, IL-2 and IL-15. **g,** Percentage of control and TIA1 TIAL1 dKO CD4 and CD8 T cells alive at day 3 after stimulation with the indicated doses of αCD3. **h,** Quantitation of the percentage of cell tracer violet dye remaining in control and TIA1 TIAL1 dKO CD4 and CD8 T cells after TCR stimulation for 3 days. In c, e, f, g and h, data from two independent experiments with at least 4 mice per genotype. In a, b and d, data pooled from three independent experiments with a minimum of 3 mice per genotype. Two-tailed Mann-Whitney tests were performed for statistical analysis.

**Supplementary Figure 3.**
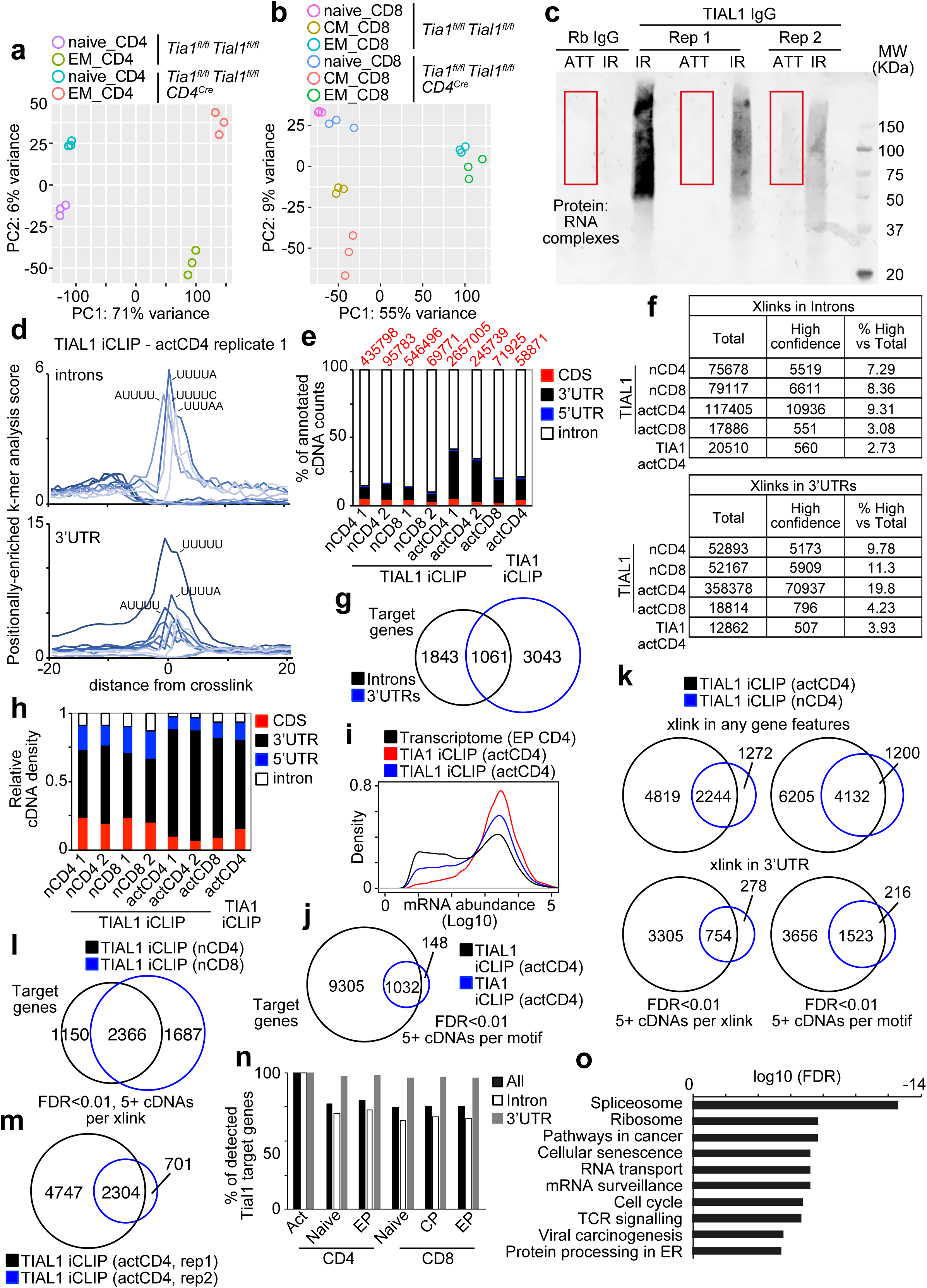
Characterization of the TIAL1 RNA interactome in T cells. a, b, Principal component analysis of the transcriptome of peripheral T cell subsets FACS sorted from the spleen of control and *Tia1^fl/fl^ Tial1^fl/fl^ CD4^Cre^* mice. **c,** Representative image showing the detection of TIAL1:RNA complexes immunoprecipitated in two independent iCLIP assays. Protein:RNA complexes were labelled with IRdye-800CW-DBCO or with the ATT-DNA linker for RNA library preparation for sequencing. An isotype antibody (rabbit IgG) was used as negative control. **d,** Positionally-enriched k-mer analysis (PEKA) in intron and 3’UTRs. Data is representative data from one of the TIA1 and TIAL1 iCLIP assays performed in naive and activated T cells. **e,** Proportion of unique cDNA counts mapped to the different mRNA transcript features in each of the iCLIP assays performed in naïve and activated T cells. Number in red indicates the number of unique cDNA counts annotated in each assay. **f,** Number of total and high-degree confidence cross-(x)link sites detected in TIA1 and TIAL1 iCLIP libraries. High-degree confidence crosslink sites were defined as being annotated by at least 5 unique cDNA counts and having an FDR<0.01. **g,** TIAL1 target genes identified in activated CD4 T cells sorted by the presence of high-degree confidence TIAL1 crosslinks in introns, 3’UTRs or both. **h,** Relative quantitation of TIA1 and TIAL1 binding density defined as the number of unique cDNA counts annotated per base of each of the indicated genomic features. **i,** Histogram showing mRNA abundance distribution of the whole transcriptome of EP CD4 T cells and, TIA1 and TIAL1 mRNA targets detected in activated CD4 T cells. **j,** Venn-diagram showing the overlap of TIA1 and TIAL1 target genes detected by iCLIP in activated T cells. The confidence degree was relaxed (5+ unique cDNA counts per binding motif defined as a window of 30 base around the crosslink) to allow annotation of TIA1 targets. **k,** Venn-diagrams comparing TIAL1 gene targets detected in naïve and activated CD4 T cells. TIAL1 target genes were defined as having at least one high confidence crosslink site in any gene feature or just in the 3’UTR. **l,** Comparison of TIAL1 gene targets with at least one high confidence crosslinking in any gene feature in naïve CD4 and CD8 T cells. **m,** Venn-diagram showing the overlap of TIAL1 gene targets detected with a high-degree of confidence in two independent replicates from activated CD4 T cells. **n,** Assessment of TIAL1 gene target expression in splenic CD4 and CD8 T cells. **o,** Pathway enrichment analysis of genes targeted by TIAL1 in CD4 T cells (WebGestalt tool, KEGG pathways).

**Supplementary Figure 4.**
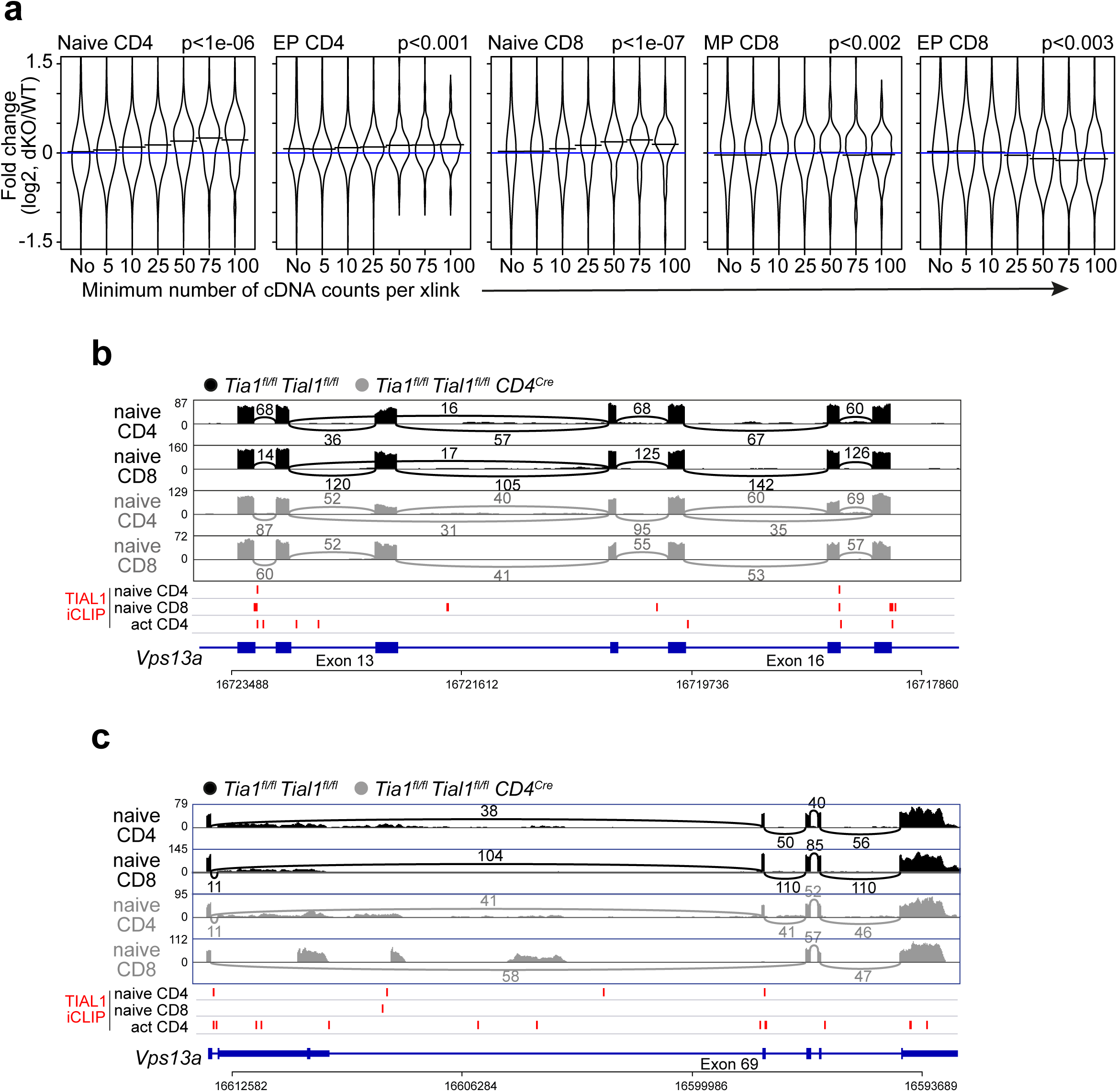
Transcriptome divergence in the absence of TIA1 and TIAL1. **a,** Global changes in mRNA expression of TIAL1 target genes in CD4 and CD8 dKO T cells compared to control T cells. Genes were grouped based on the presence of a significant crosslink site (FDR<0.05) with the indicated number of unique cDNA counts mapped per crosslink (from 5 to 100+). Kolmogorov-Smirnov tests were performed to assess for differences in the distribution of data between no targeted genes and those targets with a crosslink site in which a minimum of 50 unique cDNA counts were annotated. **b,** Sashimi plot showing the alternative splicing of *Vsp13a* exon 16 in naïve CD4 T cells from control and *Tia1^fl/fl^ Tial1^fl/fl^ CD4^Cre^* mice. **c,** Sashimi plot showing *Vsp13a* exon 69 exclusion in naïve CD8 TIA1 TIAL1 dKO T cells. In b and c, TIAL1 crosslink sites identified in naïve and *in-vitro* activated T cells are indicated in red. Values represent the number of exon-exon spanning reads annotated in each junction.

**Supplementary Figure 5.**
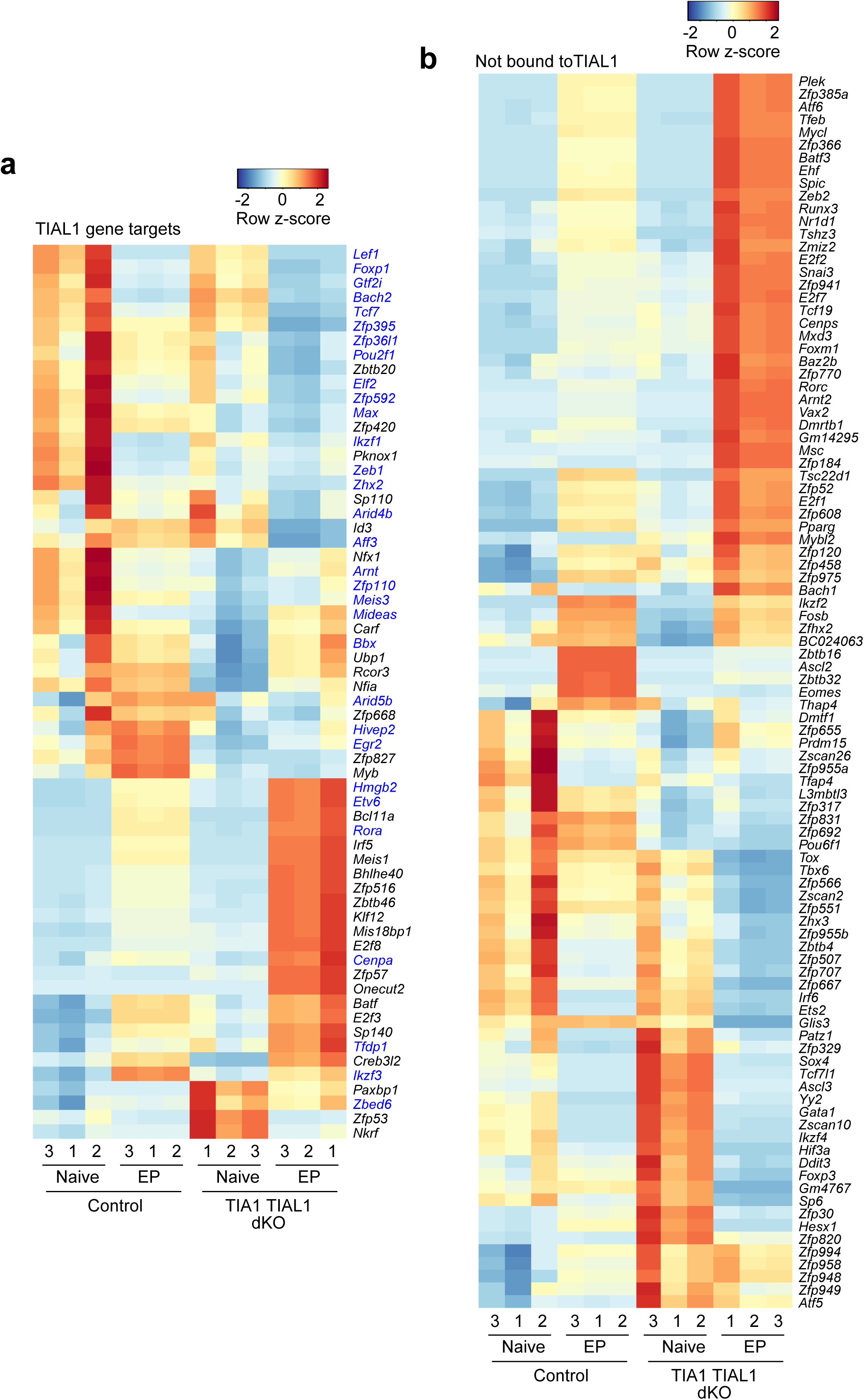
Expression of transcription factors in CD4 T cells. **a,** Heatmap showing the expression of TFs differentially expressed in TIA1 and TIAL1 dKO CD4 T cells and which transcripts are targeted by TIAL1 in the 3’UTR. In blue, TIAL1 targets annotated in naïve and *in-vitro* activated CD4 T cells. In black, TIAL1 targets only identified in activated T cells. **b,** Heatmap of TFs not targeted by TIAL1 but found differentially expressed in TIA1 and TIAL1 dKO CD4 T cells.

**Supplementary Figure 6.**
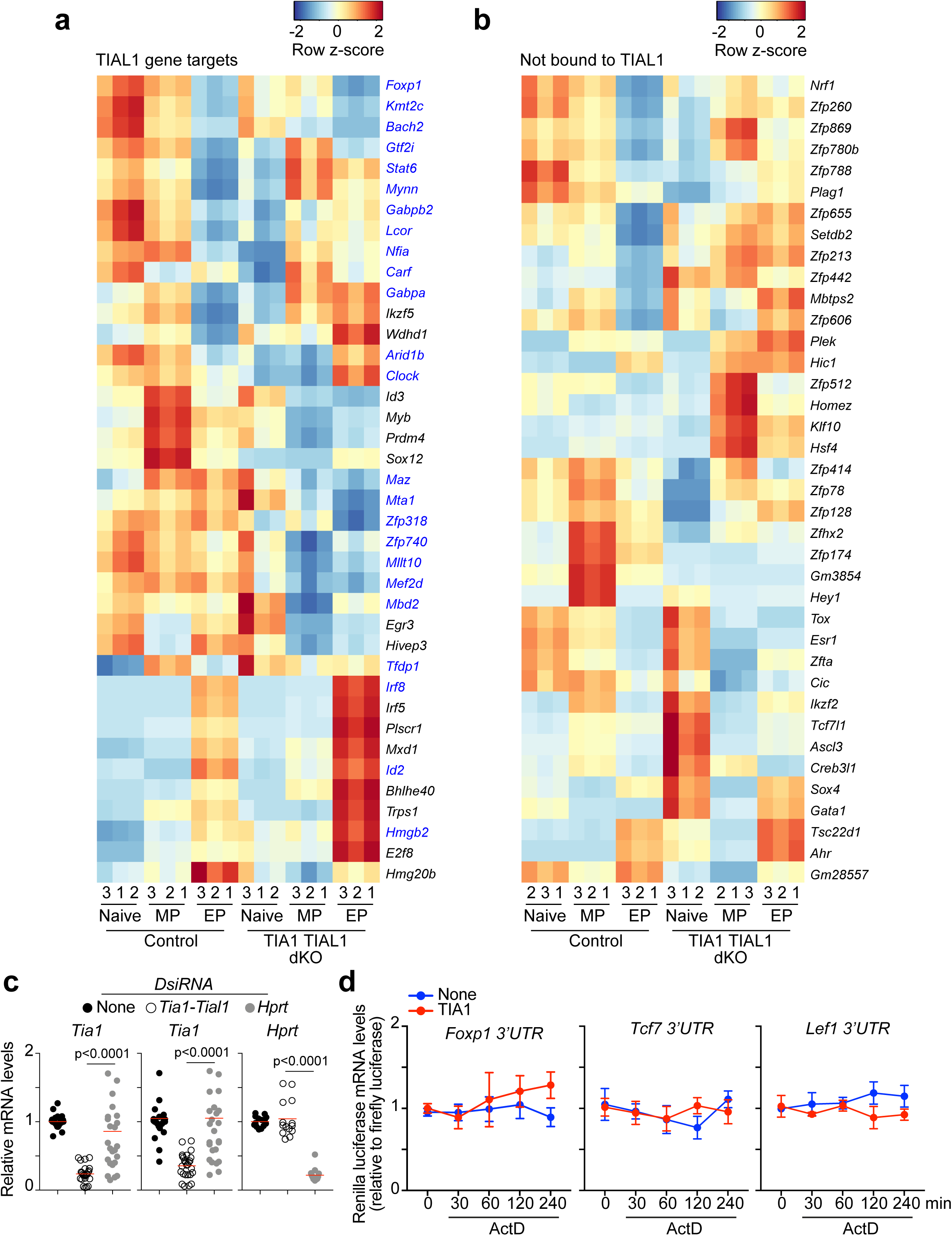
Expression of transcription factors in CD8 T cells. **a,** Heatmap showing the expression of TFs differentially expressed in TIA1 and TIAL1 dKO CD8 T cells and targeted by TIAL1. In blue, TIAL1 targets annotated in both naïve CD8 T cells and activated T cells. In black, TIAL1 targets identified only in activated T cells. **b,** Heatmap of TFs not targeted by TIAL1, but found differentially expressed in TIA1 and TIAL1 dKO CD8 T cells. **c,** DsiRNA-mediated knock down of Tia1, Tial1 and Hprt measured by RT-qPCR. Data pooled from more than five independent experiments. Two-tailed Mann-Whitney tests. **d,** Quantitation by RT-qPCR of the mRNA abundance of renilla luciferase 3’UTR reporters relative to the mRNA levels of 18S rRNA and firefly luciferase used as the internal transfection control. HEK293T cells co-transfected or not to overexpress TIA1 were treated with 5 μg/ml of ActD to block transcription at the indicated times prior cell lysis and RNA isolation using TriZol. Data pooled from two independent experiments performed by duplicate and shown as mean+SEM.

**Supplementary Figure 7.**
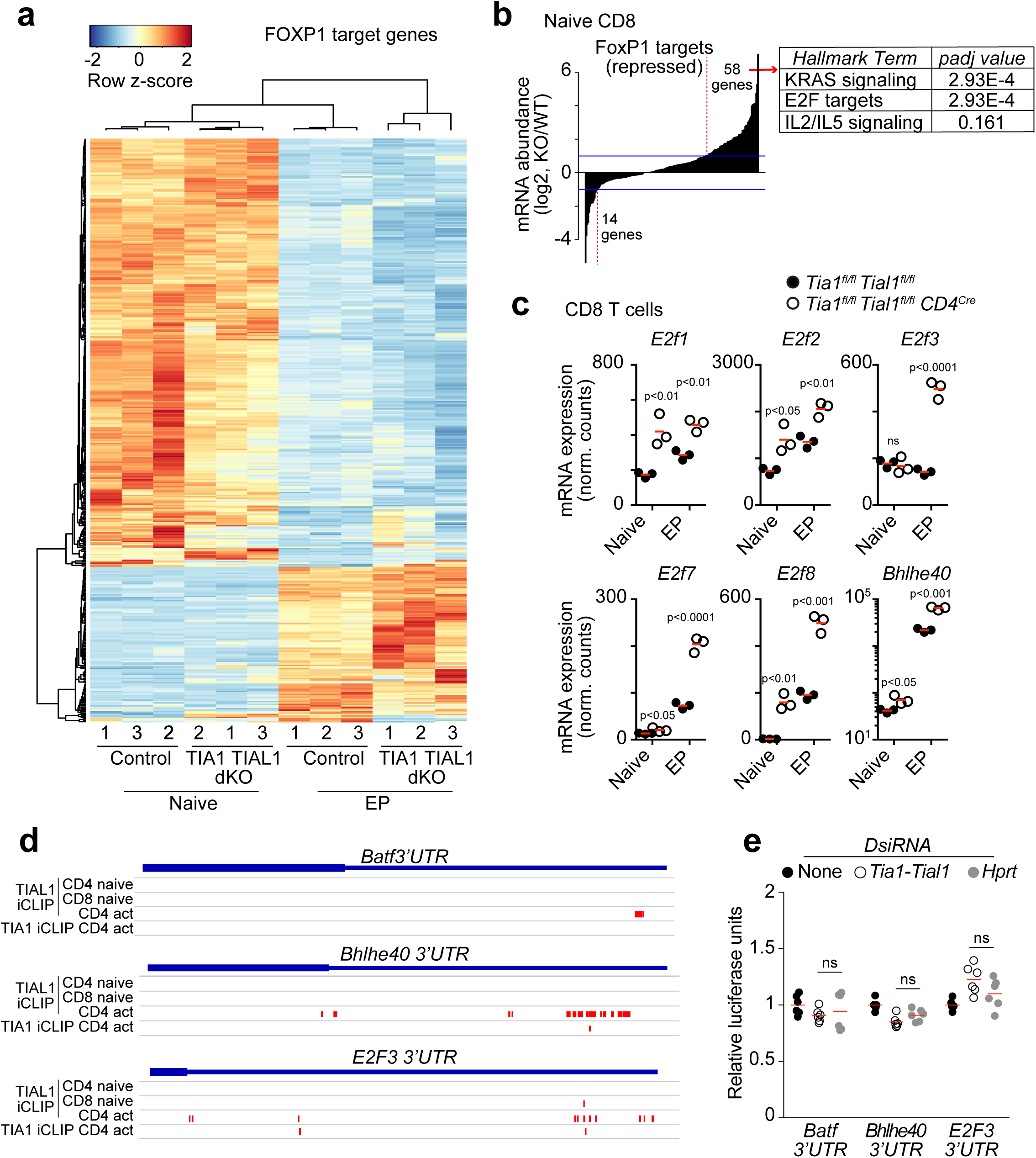
Altered expression of FOXP1 target genes in TIA1 and TIAL1 dKO CD8 T cells. **a,** Heatmap showing the expression of FOXP1 target genes in CD4 T cells. FOXP1 target genes were identified by the presence of at least one FOXP1 binding peak near or within the gene promoter (FOXP1 ChIPseq, GSE121279). Hierarchical clustering of the samples is shown. **b,** Bar plot showing the number of FOXP1-target genes with expression changes higher than 2-fold in naïve CD8 dKO T cells. Right panel, pathways enriched in upregulated FOXP1 target genes in naïve CD8 dKO T cells. **c,** Expression analysis of the indicated genes in naïve and EP CD8 T cells from control and *Tia1^fl/fl^ Tial1^fl/fl^ CD4^Cre^* mice. Mann-Whitney tests were performed for statistical analysis.

**Supplementary Table 1.** Differential expression gene analyses with DESeq2.

**Supplementary Table 2.** Annotation of TIA1 and TIAL1 RNA targets in naïve or activated T cells.

**Supplementary Table 3.** Differential expression of TIAL1 RNA targets.

**Supplementary Table 4.** Gene set enrichment analyses performed with WebGestalt.

**Supplementary Table 5.** Differential splicing analyses with rMATS.

**Supplementary Table 6.** Expression changes of FOXP1 target genes in naive CD4 T cells.

**Supplementary Table 7.** Antibodies and Reagents.

